# Endogenous insulin mediates dynamic coordination between pancreatic cancer and systemic metabolism

**DOI:** 10.64898/2026.08.26.745597

**Authors:** Jeffrey S.H. Lin, Aman A. Mohammed, Keeley G. Hewton, Jingqiang Wang, Tiffany Z.Y. Chen, Ivy S.Y. Guo, Riley S.H. Lam, Marcelo Ferraz Reinaldo, Vincent R. Richard, David F. Schaeffer, Daniel J. Renouf, Christoph H. Borchers, Seth J. Parker, Josef M. Penninger, James D. Johnson, Janel L. Kopp

## Abstract

Pancreatic ductal adenocarcinoma (PDAC) is commonly associated with obesity, diabetes, and cachexia. The pancreatic anabolic hormone, insulin, is implicated in each of these metabolic diseases, but how insulin levels affect tumor growth and relevant host physiological factors, was unclear. To address this, we transplanted orthotopic PDAC patient-derived organoids into mice consuming a hyperinsulinemia-inducing high-fat diet (HFD). We found that insulin concentrations were higher in tumors compared to circulation. Genetically reducing insulin levels reduced PDAC growth, particularly in HFD-fed mature male mice. In turn, we found the presence of pancreatic tumors increased glucose clearance and limited the expected dietary-induced gains in insulin, weight and fat mass. Interestingly, tumor growth in normal-chow-fed mice, but not HFD-fed mice, was associated with declining circulating insulin levels, as well as declines in fat and muscle mass. Together, these data provide new insights into the complex, insulin-centred interplay between PDAC and the host’s systemic metabolism that underlie cancer-associated metabolic dysfunction.

## Introduction

Pancreatic ductal adenocarcinoma (PDAC) is the most commonly diagnosed cancer of the pancreas and only 12% of patients survive five years post-diagnosis.^1,2^ This is primarily because PDAC develops silently with few symptoms and 75% of patients already have metastatic disease at diagnosis.^1,2^ Long-standing diabetes, often associated with hyperinsulinemia and obesity, increases the risk of PDAC formation.^3,4^ Additionally, the development of diabetes just prior PDAC diagnosis (new onset diabetes) is also common, with 50% of patients having diabetes at diagnosis.^5,6^ While the etiological cause of the long-term diabetes is clear, the etiological cause of the new onset diabetes is believed to be related to the destruction or disruption of the endocrine pancreatic function^6–8^ and dependence on insulin is common in these patients.^8,9^ Our previous studies demonstrated that hyperinsulinemia directly promotes initiation of precancerous lesions through insulin receptors on pancreatic acinar cells.^10^ However, because of the confounding effects of insulin on early PDAC precursor initiation, our previous work did not evaluate the role of insulin on PDAC tumors after they formed.

Insulin is produced in beta cells found in the pancreatic islets of Langerhans and then it is carried into the exocrine parenchyma by the circulatory system which then drains into the portal vein.^11^ Portal vein measurements of insulin have shown that insulin levels coming out of pancreatic circulation are higher and more variable per individual than those observed in peripheral circulation.^12^ This suggests that the whole pancreas, and not just the islet cells, are exposed to higher insulin concentrations. Insulin protein is produced from one gene in humans (*INS*) and two genes in mice (*Ins1, Ins2*). *Ins2* is the human homolog, with stronger and broader expression, while *Ins1* is a duplicated gene found only in mice and rats.^3^ In wild-type mice, two-thirds of the insulin protein is thought to be derived from the *Ins2* gene in mice.^3,13,14^ Previous studies from our laboratories have shown that genetically deleting two or three copies of *Ins1* or *Ins2* is sufficient to reduce high-fat diet (HFD)-induced weight gain without causing hyperglycemia, as well as reduce the systemic changes associated with HFD.^13,15^ We also found that deleting just *Ins2* dramatically changed precancerous lesion formation in HFD-fed mice compared to previously published papers,^16–19^ but because chow diet *Ins2* wild-type controls were not included in these studies it was unclear whether *Ins2* loss alone was sufficient to limit weight gain or reduce circulating insulin. Altogether, previous data suggested that even mild reductions in insulin production from the pancreatic beta cell can limit the extent of hyperinsulinemia, weight gain, and precancerous lesion initiation from mice prone to tumor initiation.

Diets that induce hyperinsulinemia, such as high-fat and/or high-calorie diets, can markedly influence PDAC biology and promote tumor growth.^20–24^ Consistently, insulin-lowering dietary interventions, such as low-fat, ketogenic, and caloric-restricted diets, have been shown to inhibit PDAC growth or improve treatment outcomes through various tumour-intrinsic, environmental, and inflammatory mechanisms.^25–29^ These preclinical studies primarily focused on characterizing the diet-induced changes in metabolic processes that may underlie the PDAC growth phenotypes under different dietary interventions. Many of these systemic changes coordinating the whole-body physiological responses to nutrient consumption can be downstream of insulin’s pro-anabolic and anti-catabolic functions.^30–32^ However, the effects that modulating levels of insulin specifically have on tumor growth and host metabolism were not examined.

In this study, we compared the growth of orthotopically transplanted patient-derived PDAC organoids in immunocompromised mice with or without a genetic reduction in insulin production (*Rag1^−/−^;Ins1^+/+^;Ins2^−/−^* vs *Rag1^−/−^;Ins1^+/+^;Ins2^+/+^*mice, respectively) to test the hypothesis that reducing insulin production is sufficient to lower PDAC growth. In parallel, we also gathered longitudinal changes in body weight, fasting glucose and insulin, fat mass, and lean mass, as well as glucose and insulin tolerance data at multiple time points from sham- and tumor-transplanted mice to contextualize how insulin-driven processes change with genetic reductions in insulin and in response to tumor growth. We found that *Ins2* null mice, particularly mature male mice, had modest reductions in endogenous insulin on HFD and normal chow diet (NCD) and these reductions were sufficient to reduce PDAC growth. Additionally, we found that levels of insulin were higher in the tumor interstitial fluid (TIF) as compared to serum, which were often undetectable in mice fed normal chow diet. This relative peripheral hypoinsulinemia was associated with increased catabolism in muscle and fat and potentially provided consistent micronutrient supply to the tumors. These findings highlight the complex role of insulin levels in coordinating both host nutrient homeostasis and tumor growth and provide a strong clinical rationale for monitoring and modulating insulin levels in PDAC patients.

## Results

### High Fat Diet (HFD) induces hyperinsulinemia and promotes PDO-derived tumor growth in vivo

To test our hypothesis that increasing insulin levels promote PDAC growth, we fed female and male *Rag1^−/−^* mice ^33^ a HFD (60 kcal% fat) or a normal chow diet (NCD) (18 kcal% fat) after weaning. *Rag1^−/−^* mice are expected to develop hyperinsulinemia without hyperglycemia under these conditions.^34^ We orthotopically transplanted human PDAC patient-derived organoids (PDO) derived from either a male or female patient^35^ into sex-matched mice at 8 and 12 weeks of age (male and female, respectively) to establish human PDAC in vivo (Fig 1A). We collected metabolic data at multiple time points and fluids and tissues at experimental endpoint (10 weeks post-surgery; Fig 1A). As expected, *Rag1^−/−^* mice consuming a HFD gained more weight over time than those fed a chow diet (Fig. 1B). During this time frame, fasting glucose levels fluctuated, but remained in the normoglycemic range (2-11 mM) for all *Rag1^−/−^* mice regardless of diet (Fig 1C). As expected, HFD treatment also induced a higher range of fasting circulating insulin levels and also increased HOMA-IR values compared to NCD controls in both male and female mice after 12-16 weeks of consuming HFD (Fig 1D). Consistent with the well-established effects of HFD on PDAC growth,^20–24^ the average endpoint volume and mass of pancreatic tumors in HFD-fed mice was significantly greater than those of NCD controls (Fig 1E-H). These data suggested that HFD-induced hyperinsulinemia was associated with increased growth of tumors in the pancreas.

**Figure 1.**
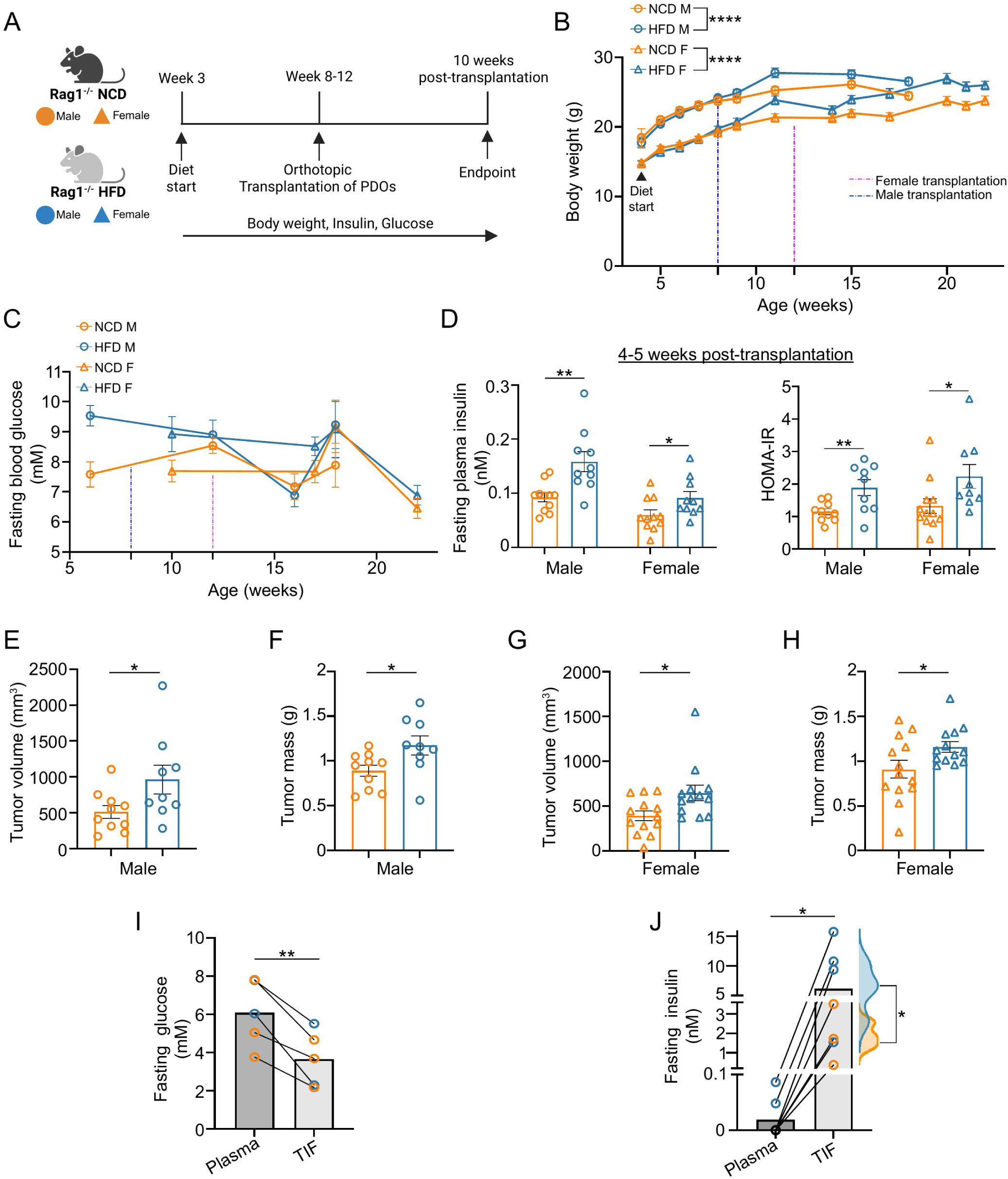
HFD promotes PDAC growth and increases insulin in tumor interstitium. (A) Schematic describing the experimental design to study orthotopic tumor progression in mice consuming a normal chow diet (NCD) or a high fat diet (HFD), n = 10-11 in each group. (B-C) Body weight (B) and fasting blood glucose (C) measurements in male (circles) and female (triangles) tumor-bearing mice, fed with HFD (blue) and NCD (orange). Values at each time point shown as mean ± SEM. ****p < 0.0001 by mixed effects test of growth rate. (D) Fasting plasma insulin and HOMA-IR values for male (5 weeks post-transplantation) and female (4 weeks post-transplantation) experimental mice under NCD (orange) or HFD (blue). Values shown as mean ± SEM *p < 0.05, **p < 0.01 by two-tailed Student t-test. (E-H) Tumor volume (E,G) and mass (F,H) from male (E-F) and female (G-H) mice at 10 weeks post-transplantation. Values shown as mean ± SEM *p < 0.05 by two-tailed Student t-test or Welch t-test. (I-J) Glucose (I) and insulin (J) concentrations present in tumor interstitial fluid (TIF) and matched plasma isolated from male mice 10 weeks post-transplantation that were fasted for 4 hours. **p < 0.01, *p < 0.05 by paired t-test. The TIF insulin values for HFD (blue) and NCD (orange) tumors were shown as half-violin plots, *p < 0.05 by Welch t-test.

### Diet-induced hyperinsulinemia increases insulin in pancreatic tumor interstitial fluid

Given the reported reductions in blood flow and the hypovascular nature of PDAC,^1,36^ as well as the reduced nutrient levels in tumor interstitial fluid (TIF),^37^ we next evaluated whether insulin was present in the tumors. We collected and analyzed glucose and insulin concentrations in TIF and the matched plasma after fasting the mice for 4 hours, ten weeks post-transplantation. We extracted TIF from 7 pancreatic tumors (3 NCD, 4 HFD) in male mice. Similar to a previous report,^37^ we found glucose was reduced in TIF relative to plasma (Fig 1I). In contrast, the fasting concentration of insulin in pancreatic TIF was at least 10 times higher than in the matched plasma (Fig 1J) for NCD-fed mice and as much as 100 times higher in TIF than plasma from HFD-fed mice (Fig 1J). These findings suggest that high insulin exposure may be a unique feature of tumors in the pancreas and systemic physiological changes that increase demand for insulin, such as those induced by HFD or high-glycemic index diets, could dramatically increase the pancreatic levels of insulin with only modest effects on circulating insulin or glucose.

### Genetically reducing insulin reduces PDAC growth in the context of HFD feeding

Given that HFD increased insulin levels in PDAC interstitium, we hypothesized that reducing the capacity for insulin production from the pancreatic islet directly would reduce HFD-accelerated PDAC growth. To test this, we crossed *Rag1^−/−^* mice with *Ins2^−/−^* mice^13,38^ to generate control mice with the full complement of *Insulin* alleles (*Rag1^−/−^;Ins1^+/+^;Ins2^+/+^*mice - hereafter referred to as *Ins2^+/+^* mice) and experimental mice with genetically fewer alleles of *Insulin* (*Rag1^−/−^;Ins1^+/+^; Ins2^−/−^* mice - hereafter referred to as *Ins2^−/−^* mice). The *Ins2* allele is predicted to produce approximately two-thirds of the endogenous insulin but its loss alone does not impair glucose homeostasis or induce overt diabetes.^3,13,14^ Using timing similar to our previous study and those of many others,^10,16–19,23,24,39,40^ we fed these mice a HFD at weaning (3 weeks of age) to promote hyperinsulinemia (Fig. S1A). We noted that 5-week-old male and female *Ins2^−/−^* mice already had a slightly smaller average body weight than controls, which persisted as the mice grew over the next 5 weeks (Fig. S1B). At 10-11 weeks of age, sex-matched human PDAC PDOs were orthotopically transplanted into both genotypes and sexes and tumor growth was assessed 10 weeks later (Fig. S1A). As expected from previous studies,^3,13,41^ the male *Ins2^+/+^* mice gained more weight than the *Ins2^−/−^*mice on HFD (Fig. S1B). *Ins2^+/+^* mice experienced some weight loss after 15 weeks of age, but this change was not as apparent for *Ins2^−/−^*mice. These initial differences in weight gain suggested *Ins2^−/−^* mice had lower insulin levels than their wild-type counterparts. At necropsy, we noted that mice with large tumors had notably smaller bodies. However, endpoint tumor volumes were comparable between *Ins2* genotypes in both the male and female cohorts (Fig. S1C-D, left panels). To further understand our necropsy observations, we correlated body weight, lean mass, or fat mass to tumor volume, we were surprised to find that they were negatively correlated (Fig. S1C-D, right panels). This negative correlation was present even in the *Ins2^−/−^* male mice and the female mice that experienced little weight loss at the end of the experiment suggesting that these negative correlations were not entirely due to cancer-induced cachexia and that body growth may be affected by tumor growth (Fig. S1B-D, right panels, figure legend). Since, IGF, which signals through similar pathways as insulin to control body growth and nutrient utilization^42,43^ is highly active in growing mice, we hypothesized that the small reductions of insulin in our model would be obscured by high IGF activity plus HFD feeding during body growth.

To examine whether insulin affects tumor growth after body growth is complete, we used EchoMRI to monitor lean mass, as a proxy of body growth, as well as fat mass in chow-fed *Ins2^+/+^* and *Ins2^−/−^* mice beginning at 5 weeks of age (Fig. S1E-H). As in the previous cohort fed HFD (Fig. S1B), both male and female *Ins2^−/−^* mice had lower body weights than controls at 5 weeks of age (Fig. S1E), driven primarily by reduced lean mass (Fig. S1F-G). Lean mass plateaued at approximately 13 weeks of age in males and 11 weeks in females, independent of genotype (Fig. S1F). In males, body weight gain was reduced in *Ins2^−/−^*mice, which was most obviously observed by discrepancies in fat mass gain (Fig. S1G). However, the relative body composition was comparable between genotypes (Fig. S1H), suggesting that insulin may also positively regulate the growth of both lean and fat mass proportionate to body weight under normal chow diet conditions.

Because the majority of our previous studies assessing HFD-induced hyperinsulinemia compared differences between three *Insulin* vs. two *Insulin* null alleles when HFD feeding started at 3 weeks of age,^13,15,18,38,40,41^ we next characterized whether insulin reduction had similar effects on mature mice by feeding them HFD for 6 weeks at 15 and 13 weeks of age for male and female mice, respectively (Fig. 2A). We measured body weight, as well as fat and lean mass (Fig 2A). We also characterized fasting insulin and glucose levels and glucose clearance (glucose tolerance test (GTT)) and insulin sensitivity (insulin tolerance test (ITT)) at multiple time points (Fig. 2A). Consistent with previous studies, mature male, and to a lesser extent female, *Ins2^+/+^* mice gained more body weight and fat mass after 6 weeks HFD feeding compared to *Ins2^−/−^*mice (Fig. 2B & 2C). Consistent with both males and females having completed their body growth, the lean mass remained stable in both males and females after the diet switch (Fig. S2A). As expected^13,34^ (Fig. 1D), switching the mice to HFD was associated with increased fasting insulin levels in both sexes, and fasting plasma insulin levels were generally higher in male *Ins2^+/+^*mice compared to females and to male *Ins2^−/−^* mice (Fig. 2D). Consistent with known *Rag1^−/−^* physiology on a HFD,^34^ fasting glucose levels remained within the normal range (Fig. 2E). Glucose intolerance is indicated by glucose levels remaining above ≥20mM 120 minutes after glucose injection in a GTT. A mild glucose intolerance (Fig. S2B-C left panels) was present in males after 6 weeks on HFD, but little to no change was observed in insulin sensitivity with HFD feeding, nor between genotypes for either metric (Fig. S2B-C right panels). The slight increases in glucose and insulin in male mice resulted in small increases in HOMA-IR and small differences were present between genotypes (Fig. S2D). In females, the average HFD-associated increases in glucose and insulin decreased to almost pre-HFD levels by 6 weeks post-diet change (Fig. 2D-E). This suggested that at least some of the females had begun to adapt to the dietary change. Altogether, these data demonstrate that reducing insulin genetically limits the effects of HFD, particularly in male mice.

**Figure 2.**
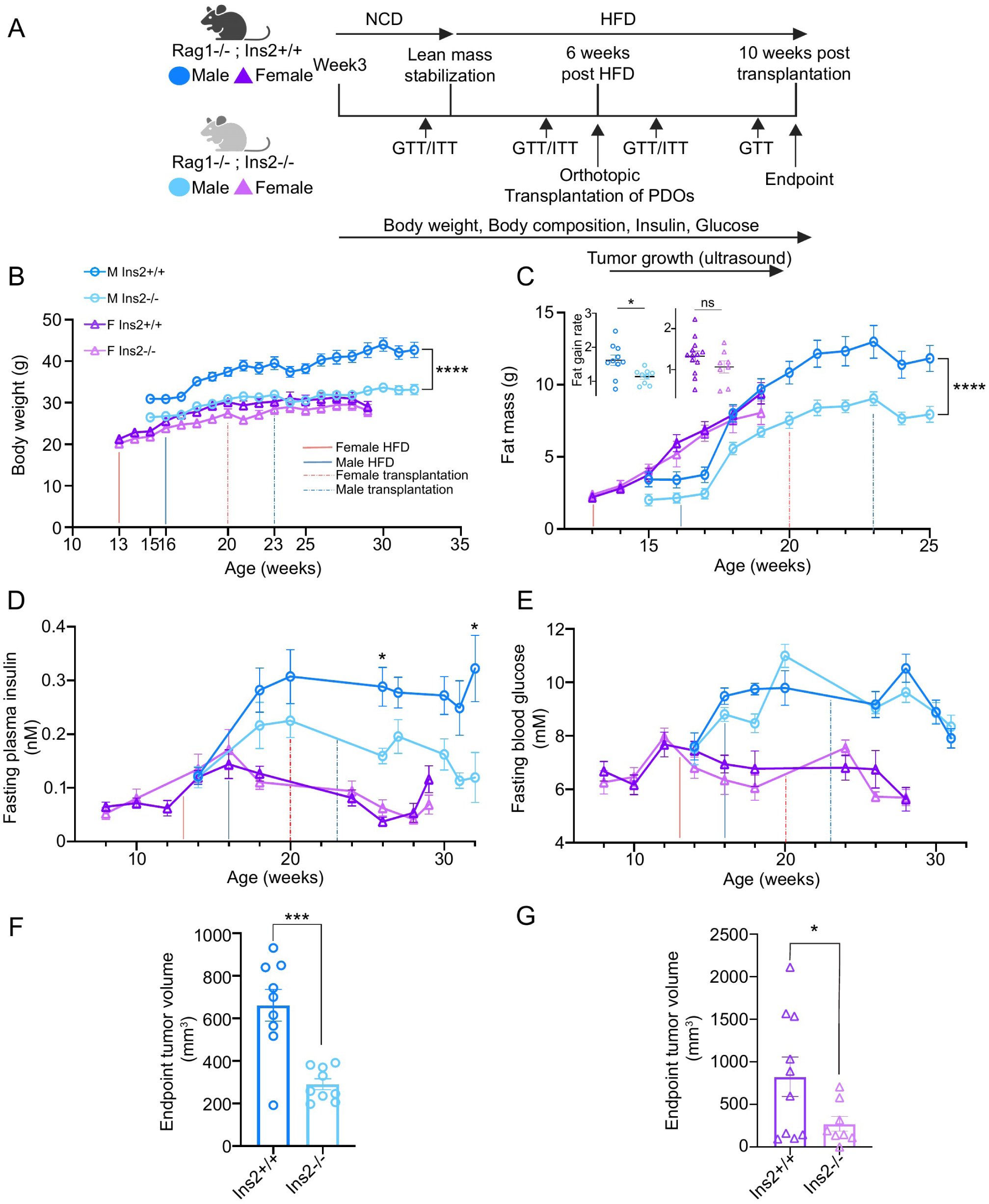
Genetic reduction of insulin reduces the effects of HFD on systemic metabolism and reduces PDAC growth. (A) Schematic describing the experimental design to study orthotopic tumor progression in *Ins2^+/+^* and *Ins2^−/−^*mice under HFD-feeding. n = 8-10 in each group. (B-C) Body weight (B) and fat mass (C) of experimental mice measured over time. The time points for HFD initiation and tumor transplantation are indicated by solid and dashed lines, respectively. Values at each time point are shown as mean ± SEM. ****p < 0.0001 by mixed-effects test of growth rate. The rate of fat gain (week 17-22 for males; week 14-19 for females) was shown in the scatter plots as mean ± SEM in (C). *p < 0.05 by Welch t-test. (D-E) Fasting plasma insulin (D) and fasting blood glucose (E) of experimental mice over time. Values at each time point are shown as mean ± SEM. *p < 0.05 by one-way ANOVA. (F-G) Calliper-measured endpoint tumor volume in male (F) and female (G) mice with varying *Ins2* doses. *p < 0.05, ***p < 0.001 by two-tailed Student t-test.

After 6 weeks on HFD, we performed PDOs orthotopic transplantation for each sex and genotype and measured tumor growth by ultrasound. The tumors were reliably detected ∼ 3-4 weeks post-transplantation and tumors in *Ins2^+/+^* mice grew larger in size than those in controls (Fig. S2E-F). The female PDO line typically grew tumors with more cystic features and was therefore often much larger in total volume than the male PDO-derived tumors (Fig. S2E-F). Consistent with this, female tumors were visibly and histologically more cystic than the male PDO-derived tumors at the experimental endpoint of 10 weeks post-transplantation (data not shown and Fig. S2G-H). Consistent with the ultrasound measurements, caliper-measured endpoint tumor volumes (Fig 2F-G), as well as tumor weights (Fig S2I-J), were on average two times greater in males and female *Ins2^+/+^* mice compared to *Ins2^−/−^* mice. These data suggested that modestly reducing the extent of HFD-induced hyperinsulinemia can significantly reduce PDAC growth in vivo.

### Pre-transplant metabolic state metrics correlate with tumor size

Serum insulin levels fluctuate throughout the day and are very individualistic,^3,38,44^ therefore a single insulin measurement is not reflective of the aggregated insulin exposure levels throughout the day. Several insulin-driven processes, like weight gain, fat storage, and HOMA-IR improve after insulin-lowering interventions are applied^13,15,38,41,45^ suggesting that these measures may be more sensitive readouts for the changes in insulin action or the insulin-regulated metabolic state. Consistent with the insulin-driven processes being sensitive records of insulin exposure, we found that an animal’s endpoint tumor volume, particularly in males, was moderately and positively correlated with body weight, fat mass, lean mass, and HOMA-IR (Extended data Table S1). However, single measures for fasting insulin or glucose did not correlate as well in males (Extended data Table S1). These correlations were mostly driven by the dynamic range of these metrics present in male *Ins2^+/+^* mice, and adding the lower values from *Ins2^−/−^* male mice to the correlation increased the power of the model to explain the data. Because multiple-time-points and/or metrics may better represent the cumulative effects of limiting endogenous insulin exposure in each mouse,^30,38^ we next integrated our longitudinal physiological measurements (Fig. 2C-E and S2A & S2D) into an insulin score for each mouse. To do this, we incorporated the average fasting insulin levels, rate of fat gain, initial lean mass, and average HOMA-IR across at least 2 time points before tumor detection or transplantation (Fig.3A-B, Extended Data Table 2, Methods) to create a single numerical score, or composite index of traits, reflecting the systemic insulin exposure within our deeply characterized cohort to which tumors are exposed. As expected, male *Ins2* knockout mice had a significantly lower average insulin score than *Ins2^+/+^*males (Fig. 3A). Additionally, the insulin score of all mice before tumors were transplanted/detected had a strong positive correlation with endpoint tumor volume in males (Fig. 3C). This correlation was primarily driven by *Ins2^+/+^* mice, suggesting that reducing the full range of insulin action in *Ins2^−/−^* mice was associated with limits on tumor size. Insulin score did not differ significantly between female genotypes (Fig. 3B) and did not correlate well with endpoint tumor size (Fig. 3D), which is consistent with the changes in these insulin score metrics being less pronounced in females.^46–48^ However, the slightly elevated status of fasting glucose and HOMA-IR in some *Ins2^+/+^*females before transplantation correlated with endpoint tumor growth (Fig 3F, Extended Data Table 2), which suggests that other readouts of the insulin-regulated metabolic state before transplantation can also be associated with tumor growth in females.

**Figure 3.**
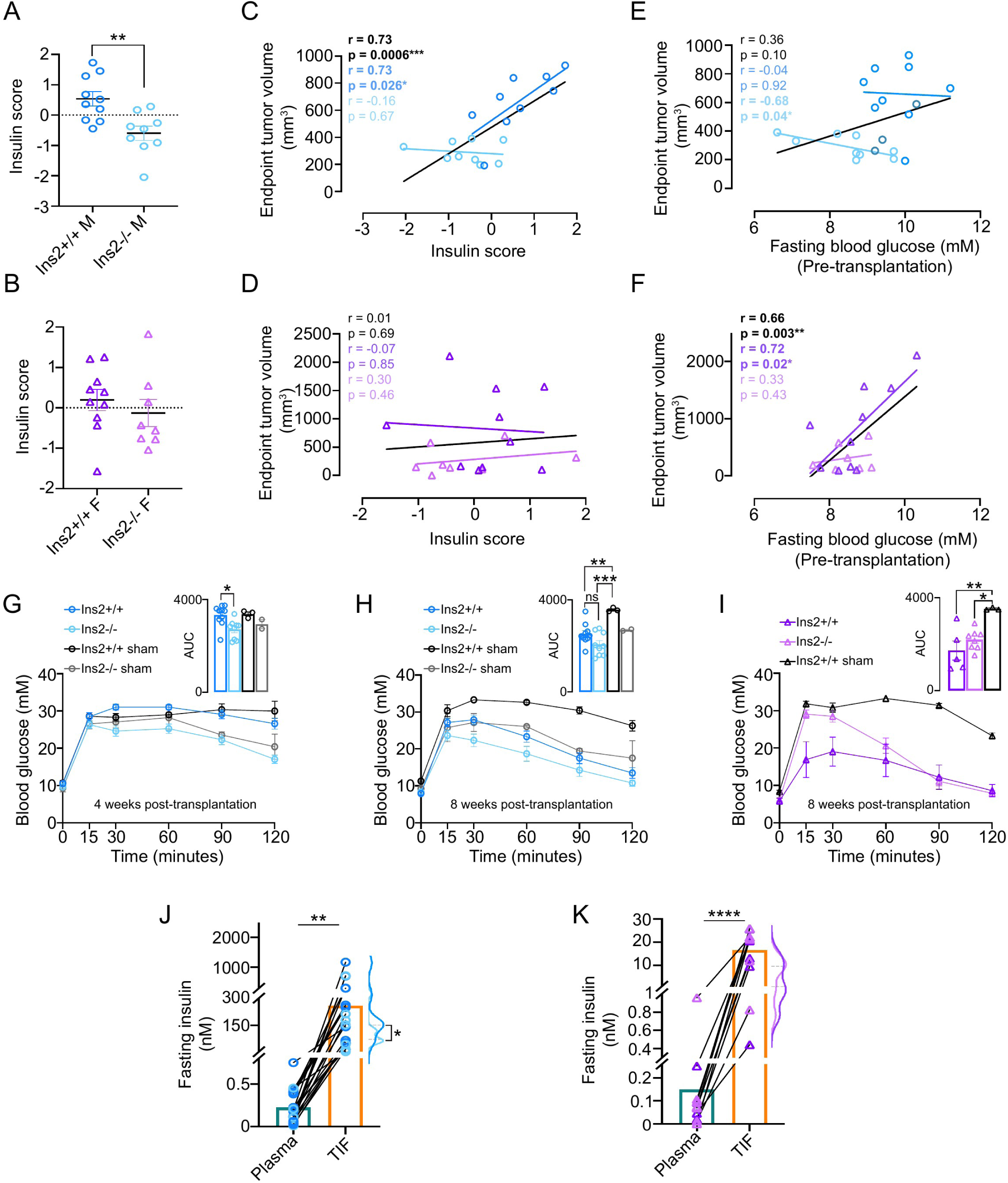
HFD induces hyperinsulinemia and high levels of insulin in TIF, but systemic readouts of hyperinsulinemia are muted by the presence of pancreatic tumors. (A-B) Insulin score estimating overall insulin exposure in HFD-fed male (A) and female (B) mice between genotypes. Values shown as mean ± SEM. **p < 0.01 by two-tailed Student t-test. (C-D) Correlation of the endpoint tumor volume with insulin score in HFD-fed male (C) and female (D) mice. r and p values were determined by Pearson correlation analysis. (E-F) Correlation of the endpoint tumor volume with the pre-transplantation fasting blood glucose in HFD-fed male (E, week 18) and female (F, week 18) mice. r and p values were determined by Pearson correlation analysis. (G-H) Intraperitoneal glucose (IPGTT) tests performed with sham- and tumor-transplanted male mice 4 (G, week 27) and 8 (H, week 31) weeks after transplantation. Area under the curve (AUC) values shown as mean ± SEM, *p < 0.05, **p < 0.01, ***p < 0.001 by one-way ANOVA. (I) IPGTT tests performed with sham- and tumor-transplanted female mice 8 weeks after transplantation (week 28). AUC values shown as mean ± SEM. (J-K) Bar plots showing the insulin concentrations in TIF and the matched plasma at fasting in HFD-fed male (J) and female (K) mice, **p < 0.01, ****p < 0.0001 by paired t-test. The average TIF insulin values in tumor from *Ins2^+/+^* and *Ins2^−/−^* hosts were shown as half-violin plots, *p < 0.05 by Welch t-test.

### Hyperinsulinemia and its associated sequelae is curtailed in mature tumor-transplanted mice

Previous studies have suggested that these insulin-regulated metabolic metrics can be affected by the presence of cancer^5,6,49,50^ and approximately 50% of PDAC patients have dysglycemia,^5,6^ therefore we next examined whether these insulin-related physiological measures changed in mice as tumors grew (Fig. 2A). To get a baseline for the expected HFD-induced responses of this cohort, we monitored the weight gain, glucose and insulin levels, as well as performed ITTs and GTTs in a small number of sham-transplanted *Ins2^+/+^*and *Ins2^−/−^* mice from the same mating cohort. Lean and fat mass were also assessed in sham and a limited number of tumor-transplanted mice after transplantation. As expected, surgery caused a transient decrease in weight for all mice (Fig 2B), which appeared to be primarily due to transient loss of fat mass (Fig 2C-appropriate time only assessed in male mice). After recovery, sham-transplanted male *Ins2^+/+^* mice continued to gain more weight and fat mass (Fig. S3A-C) when compared to sham-transplanted *Ins2^−/−^* male mice and tumor-transplanted mice of both genotypes (Fig. S3A-C). Sham- and PDO-transplanted *Ins2^+/+^* female mice, in contrast, had little to no increases in weight after recovering from surgery (Fig 2B & S3A-C). Weight loss indicative of cancer-associated cachexia (>10%) was not present by the experimental endpoint. Together, these data suggested that presence of pancreatic tumors reduced the expected HFD-induced weight gain in male mice with full insulin gene dosage.

Next, we examined whether glucose homeostasis changed with tumor transplantation. Post-transplantation, we observed that fasting levels of glucose varied, but remained within upper- and mid-normoglycemic range for tumor- and sham-transplanted male and female mice, respectively (Fig S3D). We also assessed the response of sham- and tumor-bearing mice to a glucose bolus at 4 and/or 8 weeks after transplantation (Fig. 3G-I). Consistent with the long-term effects of HFD on male mice,^40,51,52^ the slight glucose intolerance observed in 21-week-old HFD-fed male mice before transplantation (Fig. S2C) worsened after a further 6 weeks on HFD (4 weeks post-transplantation-Fig. 3G). Glucose clearance was better in *Ins2^−/−^* mice compared to *Ins2^+/+^* mice; and sham- and tumor-transplanted mice were similar within genotypes (Fig. 3G). This genotypic difference in glucose clearance persisted in sham-transplanted mice 8 weeks after surgery, however tumor-transplanted mice had significantly improved glucose clearance (Fig.3H-I). Interestingly, tumors were already measurably larger in *Ins2^+/+^* mice compared to *Ins2^−/−^* mice at this time point (Fig. S2E) and *Ins2^+/+^* tumor-transplanted males had more improvement in glucose clearance (Fig. 3H). Female mice showed better glucose clearance than males after 6 weeks on HFD (Fig. S2C). By 8 weeks after surgery, sham-transplanted *Ins2^+/+^* females had also developed glucose intolerance (Fig. 3I, *Ins2^−/−^* females were not evaluated). Similarly to males, tumor-transplanted females had better glucose clearance at 8 weeks post-transplantation when compared to sham-transplanted females (Fig. 3I). Together, these data suggested that the HFD-mediated effects on glucose homeostasis were improved coincident with increasing pancreatic tumor size in mice.

To determine whether the reduced effect of HFD on tumor-bearing mice was associated with reductions in hyperinsulinemia and insulin resistance, we measured fasting insulin at multiple time points post-transplantation, and also assessed insulin resistance with ITT and HOMA-IR calculations (Fig. 2A). Although sham-transplanted *Ins2^+/+^* females become glucose intolerant by 28 weeks of age (Fig. 3I), both tumor-transplanted and sham-transplanted females slowly lowered their levels of insulin to pre-HFD levels by 28 weeks of age and maintained low HOMA-IR levels (Fig S3G). This is consistent with previous studies^48,53^ indicating that females maintain better insulin sensitivity. Male Bl6 mice, in contrast, more frequently become insulin resistant and dramatically increase their circulating insulin levels with age.^10,40,52–55^ Indeed, we found that fasting plasma insulin and HOMA-IR dramatically increased in sham-transplanted *Ins2^+/+^* and *Ins2^−/−^* male mice, particularly by 30 weeks of age (Fig. S3E & F). These increases were generally less in *Ins2^−/−^* male mice compared to controls, suggesting that they may have better insulin sensitivity. However, the response to insulin during an ITT at 4 weeks post-transplantation (Fig. S3G, only tumor-transplanted males were tested) was normal and similar between genotypes, suggesting, at least at this time point, the mice had not yet developed overt insulin intolerance. In contrast to the dramatic increases in insulin and HOMA-IR in sham-transplanted mice, tumor-transplanted male *Ins2^+/+^* and *Ins2^−/−^* male mice experienced little to no change in these metrics after transplantation (Fig. 2D, S2D and S3E & F). These data strongly suggest that the presence of a tumor in the pancreas blunts the degree of hyperinsulinemia and hyperinsulinemia-driven processes expected when mice consume HFD.

### Insulin levels are extremely high in pancreatic tumors from mature mice consuming HFD

Although HFD-diet induced hyperinsulinemia was blunted in tumor-bearing mice, our previous studies of TIF suggested that HFD induced higher pancreatic insulin levels when compared to serum levels (Fig. 1J). In this mature HFD cohort, where HOMA-IR levels were dramatically higher than in the young cohort (Fig. 1D vs S2D), we found that TIF fasting insulin levels were between ∼5 and 1000 nM and 0.4-26 nM from mature males and females, respectively, compared to the 0.03-0.8 nM concentrations found in the matched fasting serum (Fig. 3J-K). In contrast, fasting TIF glucose levels were lower than serum (Fig. S3H-I), as previously reported (Fig. 1J).^37^ This suggested that serum levels of insulin are not entirely reflective of the extent of pancreatic insulin and that high insulin levels could be constitutively present in pancreatic tumors in this context. Notably, the average TIF insulin concentration was significantly lower in the *Ins2^−/−^*males than in wild-type controls (Fig. 3J), indicating that genetically reducing *Insulin* reduces exposure of pancreatic cells to insulin. Altogether, these data suggest that pancreatic tumors in mice consuming a HFD can be exposed to high insulin levels even though the systemic effects of hyperinsulinemia are masked.

### Modest reductions in insulin in chow-fed mice reduce PDAC growth

To test whether the extremely high pancreatic insulin levels induced by HFD were necessary for insulin to promote PDAC growth, we fed *Ins2^+/+^*and *Ins2^−/−^* mice with only NCD and transplanted the same PDOs in a sex-matched manner at time points matched to the HFD experiments (Fig. 4A). As before, we collected multiple physiological measurements over the course of the experiment and measured tumor growth by ultrasound (Fig. 4A). As observed previously, *Ins2^+/+^*and *Ins2^−/−^* mice again had differences in body weight and lean mass at 15 weeks of age (Fig 2B, S2A, Fig. 4B, S4A). Males further increased in weight by about 5% regardless of genotype over the next few weeks before transplantation, but weight remained stable in females (Fig. 4B). The males gained primarily fat mass and not lean mass during this period (Fig. 4C, S4A). We also found that fasting glucose levels were variable, but in the normal range before transplantation (Fig. 4D); and GTT and ITT responses were similar between genotypes (Fig. S4B). Fasting insulin levels were low and variable, and a modest difference in insulin was present between genotypes for tumor-transplanted male, but not female, mice (Fig. 4E). Consistent with these data, HOMA-IR was variable, but also not different between genotypes (Fig. S4C). Even though changes in insulin levels were minor, we still found that PDO-derived tumors in male *Ins2^+/+^* mice were larger than those from *Ins2^−/−^* male mice (Fig. 4F, S4D-E), albeit with a smaller difference than observed under HFD conditions. Initially, tumors in female *Ins2^−/−^*mice appear to grow larger than those in *Ins2^+/+^* mice (Fig. S4F), but measurements by caliper and weight showed no difference at experimental endpoint (Fig. 4G and S4G). Potentially consistent with the cystic nature of the tumors derived from the female PDO line, these tumors tended to be larger in size than the male-derived PDO line (Fig. 4F-G). But no consistent difference in the extent of cysts nor histology between genotypes was observed for males or female tumors (Fig. S4H-I). Collectively, these results indicate that a modest reduction in the endogenous insulin levels was sufficient to restrain PDAC growth, particularly in male mice, even in the absence of the extreme levels of diet-induced hyperinsulinemia.

**Figure 4.**
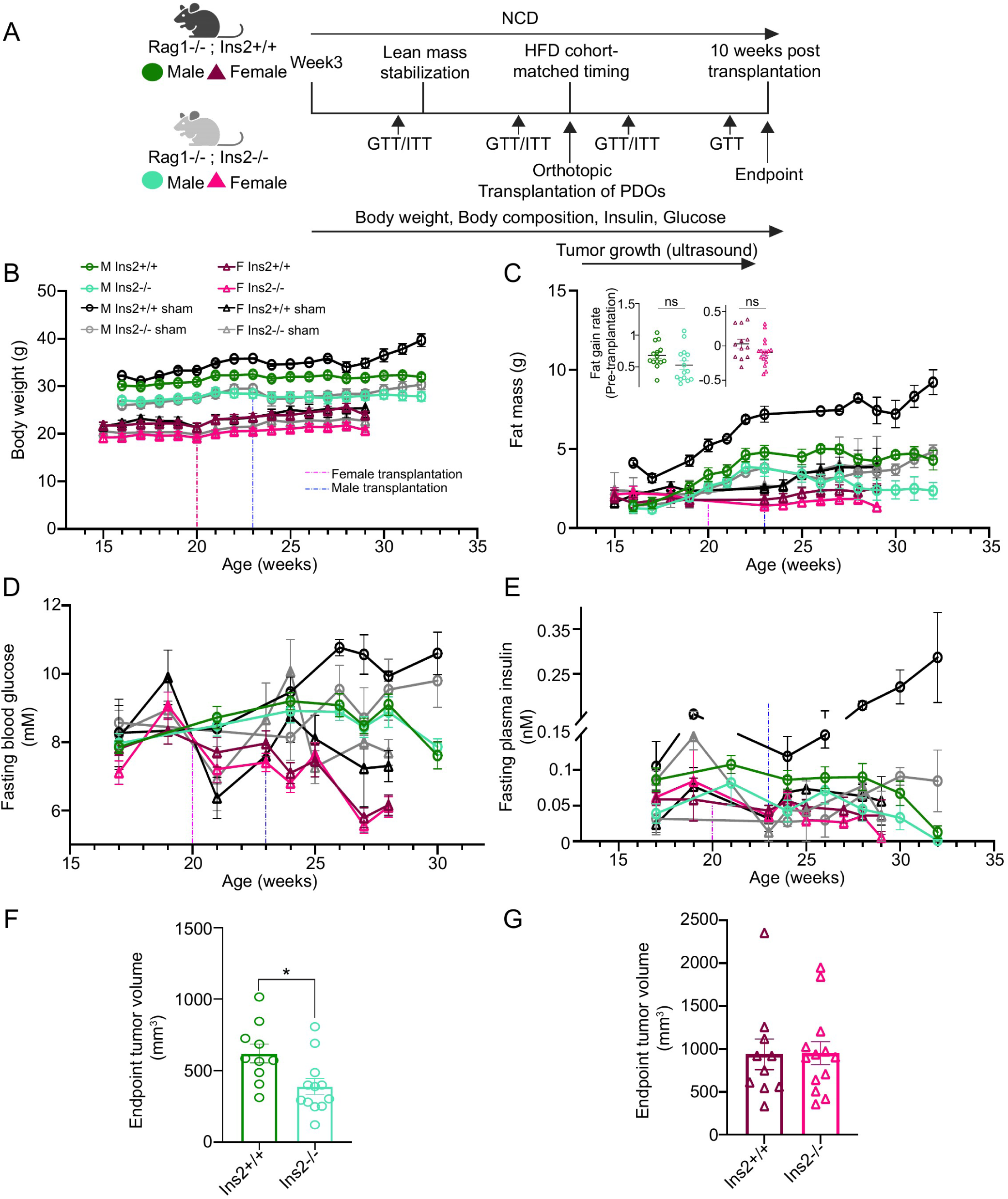
Modest reductions of insulin in normal chow diet mice suppresses PDAC growth. (A) Schematic describing the experimental design to study orthotopic tumor progression in *Ins2^+/+^* and *Ins2^−/−^* mice under chow-feeding. n =10-13 in each tumor-transplanted group; n = 3 in each sham-transplanted group. (B-C) Body weight (B) and fat mass (C) of experimental mice measured over time. The time points for tumor transplantation are indicated by dashed lines. Values at each time point are shown as mean ± SEM. The rate of pre-transplantation fat gain used for insulin score calculation (week 16-22 for males; week 14-19 for females) was shown in the scatter plots as mean ± SEM in (C). After transplantation, fat mass values of tumor-transplanted mice were obtained from 3 animals (n=3 for each group of tumor-transplanted mice). Dashed line indicates sham or tumor transplantation. (D-E) Fasting blood glucose (D) and fasting plasma insulin (E) of experimental mice over time. Values at each time point are shown as mean ± SEM. Dashed line indicates sham or tumor transplantation. (F-G) Calliper-measured endpoint tumor volume in chow-fed male (F) and female (G) mice with varying *Ins2* doses. Values shown as mean ± SEM, *p < 0.05 by two-tailed Student t-test.

### *Insulin* reduction on normal chow diet creates metabolic states associated with tumor size

Since pre-transplantation levels of insulin function correlated with endpoint tumor volume in male mice fed HFD (Fig. 3C), we next examined whether this relationship also existed in the absence of overt hyperinsulinemia. To do this, we used the collected physiological measurements for each mouse to calculate correlations with tumor size, as well as a composite insulin score similar to our previous calculations (Extended data table S1-2; Methods). We found that *Ins2^+/+^*male mice again had a higher average insulin score compared to *Ins2^−/−^*mice (Fig. 5A). These insulin scores correlated well with endpoint tumor volume for male mice regardless of genotype and when the range of insulin/tumor growth was considered across genotypes this relationship was significant (Fig. 5B). In contrast, female mice predominantly had similar insulin scores (Fig 5C) and no correlation was present between tumor volume and insulin score before transplantation (Fig. 5D). Consistent with female mice having very little response to *Insulin* reduction on normal chow diet before transplantation, the fasting glucose prior to transplantation also did not correlate with tumor growth in females on chow diet (Extended data table S1). This suggests that, at least for males transplanted with PDOs, if a higher range of insulin activity is present before tumor transplantation, tumor sizes will likely reflect those differences.

**Figure 5.**
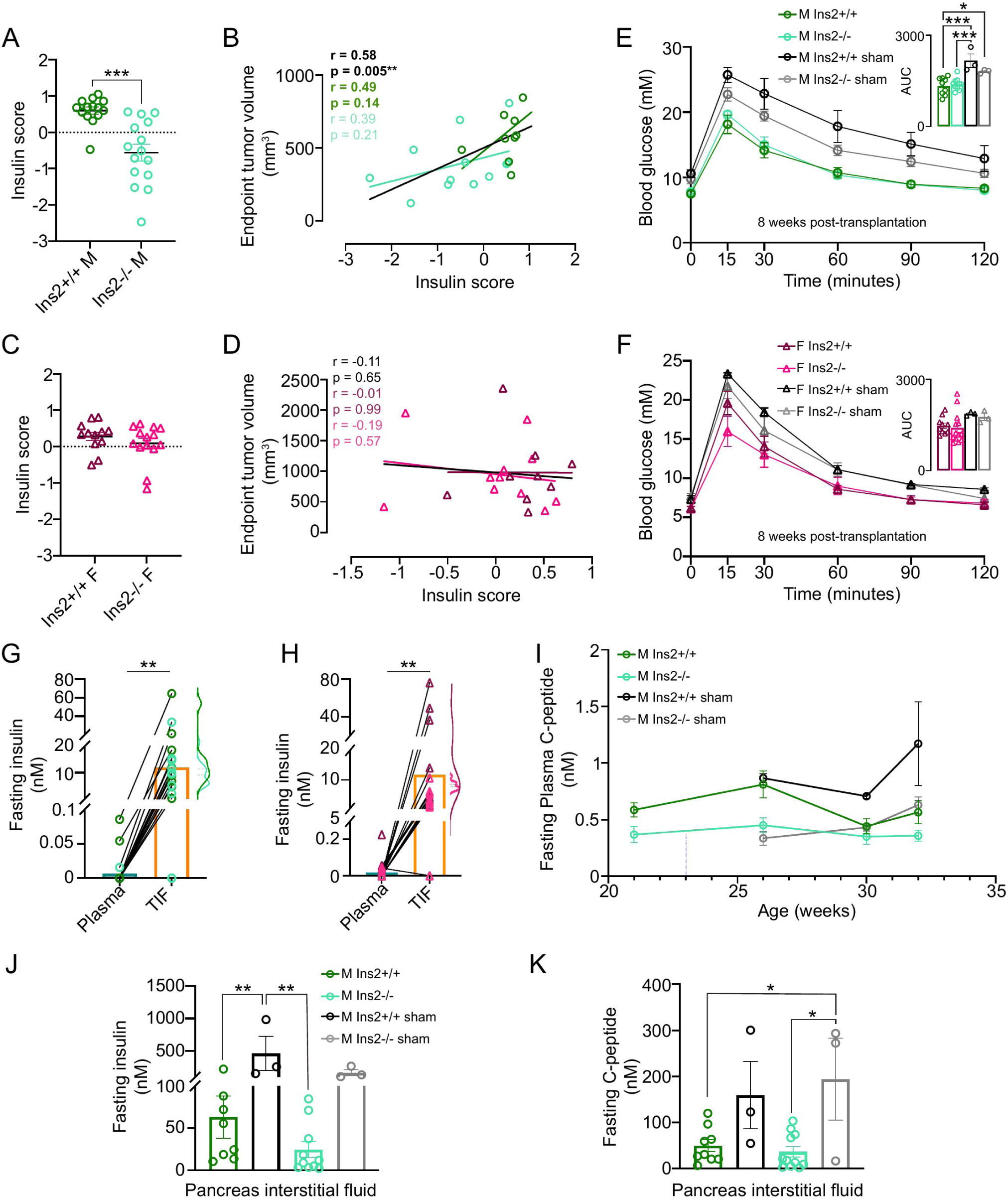
Glucose clearance improves coincident with reductions in peripheral and pancreatic insulin levels in tumor bearing mice. (A) Insulin score estimating overall insulin exposure in chow-fed male mice. Values shown as mean ± SEM, ***p < 0.001 by Welch t-test. (B) Correlation of the endpoint tumor volume with insulin score in chow-fed male mice. r and p values were determined by Pearson correlation analysis. (C) Insulin score estimating overall insulin exposure in chow-fed female mice. Values shown as mean ± SEM. (D) Correlation of the endpoint tumor volume with insulin score in chow-fed female mice. r and p values were determined by Pearson correlation analysis. (E-F) Intraperitoneal glucose (IPGTT) tests performed with sham- and tumor-transplanted male (E, week 31) and female (F, week 28) mice 8 weeks after transplantation. Area under the curve (AUC) values shown as mean ± SEM, *p < 0.05, ***p < 0.001 by one-way ANOVA. (G-H) Bar plots showing the insulin concentrations in TIF and the matched plasma at fasting in chow-fed male (G) and female (H) mice, **p < 0.01 by paired t-test. The average TIF insulin values in tumor from *Ins2^+/+^*and *Ins2^−/−^* hosts were shown as half-violin plots. (I) Fasting plasma C-peptide of chow-fed male over time. Dashed line indicates sham or tumor transplantation. Values at each time point are shown as mean ± SEM. (J-K) Concentrations of insulin (J) and C-peptide (K) in pancreas interstitial fluid samples isolated from the normal pancreas tissue of sham- and tumor-transplanted male mice. Values shown as mean ± SEM, *p < 0.05, **p < 0.01 by one-way ANOVA.

### Tumor-transplanted NCD-fed mice developed peripheral hypoinsulinemia associated with increased fat and muscle catabolism as tumors grew

Given that tumors affected the expected insulin-driven processes in the context of a HFD, we next examined how tumors affected insulin and its downstream actions when excess nutrients were not present. Both sham- and tumor-transplanted female mice gained about 10% more weight after recovering from transplantation (week 21-24) and experienced normal variances in weight thereafter (Fig. 4B). Because these weight increases also occurred in sham-transplanted females (Fig. 4B), these changes were likely due to a normal period of weight gain in mature female mice. Interestingly, although weight gain did not differ between sham- and tumor-transplanted female mice of similar genotype, the tumor-transplanted females appeared to gain no fat mass after 24 weeks of age while sham-transplanted mice gained a small amount of fat mass (Fig. 4C and S4A). Indeed, gonadal and subcutaneous fat depots were significantly lighter in tumor-transplanted mice at endpoint compared to sham-transplanted even when normalized to body weight (Fig. S5A). We also weighed skeletal muscle, heart, and liver and found that only skeletal muscle mass was smaller in tumor-bearing mice, and the liver increased as a percentage of total body weight (Fig. S5B-C). This suggested that in tumor-transplanted females, increasing tumor and liver weight could offset the absence of fat gain and loss of muscle mass to result in little change in total body weight and lean mass (Fig. 4B & S4C). Because these changes were very similar regardless of genotype, this suggested that tumor-transplanted females of both genotypes may have increased peripheral catabolism.

In contrast to females, sham-transplanted males, particularly *Ins2^+/+^*males, began to gain dramatically more body weight, as well as fat mass around 28 weeks of age (Fig. 4B-C and S4A). *Ins2^−/−^* mice had a less pronounced change in weight and fat mass beginning at 30 weeks of age (Fig. 4B-C and Fig. S4A) suggesting that reduced *insulin* gene dosage affected weight gain in NCD-fed males. Tumor-transplanted male mice of both genotypes did not gain weight or fat mass after recovering from transplantation and had similar body weight and fat mass as sham-transplanted *Ins2^−/−^* mice (Fig. 4B). This suggested that the presence of tumors in wild-type mice can phenocopy the reduced insulin action associated with the absence of *Ins2*. Although body weight did not drop dramatically in tumor-transplanted *Ins2^−/−^* mice, these mice steadily lost lean and fat mass beginning 3 weeks after transplantation (Fig. 4C and S4A). This suggested that the presence of pancreatic tumors in mice with reduced insulin production capacity was associated with increased catabolism. Confirming these observations, we found that relative fat depots weights in males of both genotypes were less than in sham-transplanted animals (Fig. S5D). Additionally, the difference in absolute muscle weight was greater by genotype in tumor-transplanted compared to sham-transplanted male mice (Fig. S5E). In contrast to female mice, relative liver weights in sham- and tumor-transplanted males were similar (Fig. S5F). Altogether, these data suggested that the expected gain in body weight or fat mass did not occur in the tumor-transplanted NCD-fed mice, and additional losses of muscle and fat may have occurred in mice with low insulin.

Mobilization of nutrients from fat and muscle is expected to indirectly increase circulating glucose levels.^56,57^ However, we found that fasting glucose levels tended to be lower 3-5 weeks before experimental endpoint in tumor-transplanted mice compared to sham-transplanted mice (Fig. 4D). This suggested that the presence of the tumor affected the levels of glucose present during fasting. We next evaluated insulin sensitivity and glucose clearance. At 4 weeks post-transplantation, GTT responses for males and females were similar between genotypes and between sham- and tumor-transplanted mice (Fig. S5G-H). After 8 weeks of tumor growth, tumor-transplanted mice of both sexes tended to have slightly better glucose clearance than the sham-transplanted counterparts (Fig. 5E-F). This suggested that the presence of pancreatic tumors reduced glucose levels by improving clearance regardless of genotype or sex.

Insulin is the primary anabolic hormone in the body,^30,32,58^ and catabolism in the fat and muscle tissues normally occurs when insulin levels are low.^13,59,60^ Therefore, we next examined whether insulin levels were reduced in tumor-transplanted NCD-fed mice compared to sham controls. Consistent with the increased weight in sham-transplanted *Ins2^+/+^*male mice (Fig. 4B), fasting levels of insulin and HOMA-IR began to steadily increase after recovery from surgery compared to all other groups (Fig. 4E and S4C). After tumor transplantation, insulin levels remained stable with a consistent difference seen between genotypes for male mice until 1-3 weeks before the experimental endpoint (Fig. 4E). HOMA-IR values fluctuated, but remained low and were similar between genotypes after transplantation (Fig. S4C). This suggested that the presence of tumors in the pancreas limited the normal increases in insulin for males in the *Rag1^−/−^*Bl6 background. Seven to nine weeks after tumor transplantation, however, we observed that some mice began to have undetectable levels of fasting insulin using an ELISA assay that should be able to detect 0.005 nM levels of insulin (Extended data table 3, Fig. 4E). At experimental endpoint, 87% and 78% of all tumor-transplanted male and female mice, respectively, had undetectable levels of insulin in fasted serum (Fig. 4E, 5G-H and Extended data table 3). Interestingly, both *Ins2^−/−^*male mice and both female genotypes more often had undetectable insulin levels in the blood and that frequency increased towards the endpoint (Extended data table 3). This suggested that female mice and male mice with reduced insulin may not have had enough insulin induce support anabolism in peripheral tissues when tumors were present.

In contrast to the serum, insulin levels were high in the TIF (Fig. 5G-H). This suggested that insulin was still being made and tumors were still exposed to these higher levels. Consistent with insulin still being produced, 1) C-peptide levels in the serum were still present at levels similar to pre-transplantation levels (Fig. 5I); 2) interstitial fluid isolated from the non-transplanted portions of the same pancreas also had higher insulin levels than serum (Fig. 5G & 5J); and 3) the mice were not hyperglycemic (Fig. 4D). Although insulin was still being made in tumor-transplanted mice, the pancreatic interstitial fluid (pancIF) levels of insulin and C-peptide levels in tumor-transplanted mice were generally lower than in pancIF collected from the sham-transplanted mice (Fig. 5J-K). Notably, the pancIF insulin levels reflected the difference in *insulin* gene dosage (Fig. 5J). Altogether, these data suggest that the presence of orthotopic PDO-transplanted tumors can reduce the pancreatic levels of insulin, but there is still more insulin present in the tumor than in circulation at experimental endpoint. This relative lack in circulation is associated with reductions in lean and fat mass in mice. This suggested that conditions which promote increased insulin production from the islet, such as HFD, could alleviate this peripheral lack of insulin. Indeed, HFD-fed tumor-transplanted female and male mice maintained fasting serum insulin levels, body weights, lean masses, and fat masses similar to that observed at the time of transplantation throughout the 10 weeks (Fig. 2A-B & S2A). However, TIF levels, and likely pancIF levels became excessively high (Fig. 3J-K). In addition, the excessive glucose levels and insulin resistance that should have been present due to the HFD did not develop likely because the excess glucose was cleared quickly and likely used by the tumor.

### *Insulin* gene dosage regulates PDAC cell proliferation

Given that changes in insulin function in the body are associated with differences in tumor growth and insulin levels are still high in the TIF at the experimental endpoint, we used endpoint-collected bulk tumor samples to assess the proteomic and metabolomic changes created by *Insulin* gene reduction in male mice. We focused on male mice because the effect of *Insulin* gene dosage on tumor growth was consistent in this sex across both diet conditions. Specifically, we performed LC-MS for polar metabolites, including absolute quantification of glucose and amino acids, in TIF and plasma from control and *Ins2^−/−^* male mice under both diet conditions. We also performed unbiased data-independent acquisition (DIA) proteomic analysis by LC-MS/MS using a randomly selected piece of solid tumor from *Ins2^+/+^* and *Ins2^−/−^* male mice under both diet conditions. We measured the relative abundance of 187 metabolites in TIF and plasma from control and *Ins2^−/−^* male mice under both diet conditions by LC-MS (Extended Data Table 4). As expected,^37^ the isolated TIF generally contained relatively lower levels of glucose, arginine, and tryptophan compared to matched plasma, whereas lactate, alanine, glutamate, and aspartate were generally higher in TIF (Fig. S6A). However, we observed only sample type clustering of the metabolites and no significant differences were found between genotypes or diets (Fig 6A). Furthermore, there appeared to be a negative concordance between genotypes across diets in the TIF metabolite levels (Fig. S6B). These data suggested that the TIF metabolite changes were an indirect compensatory effect of the animal to the different genotypes and diets and not a consistent readout of insulin-driven changes.

**Figure 6.**
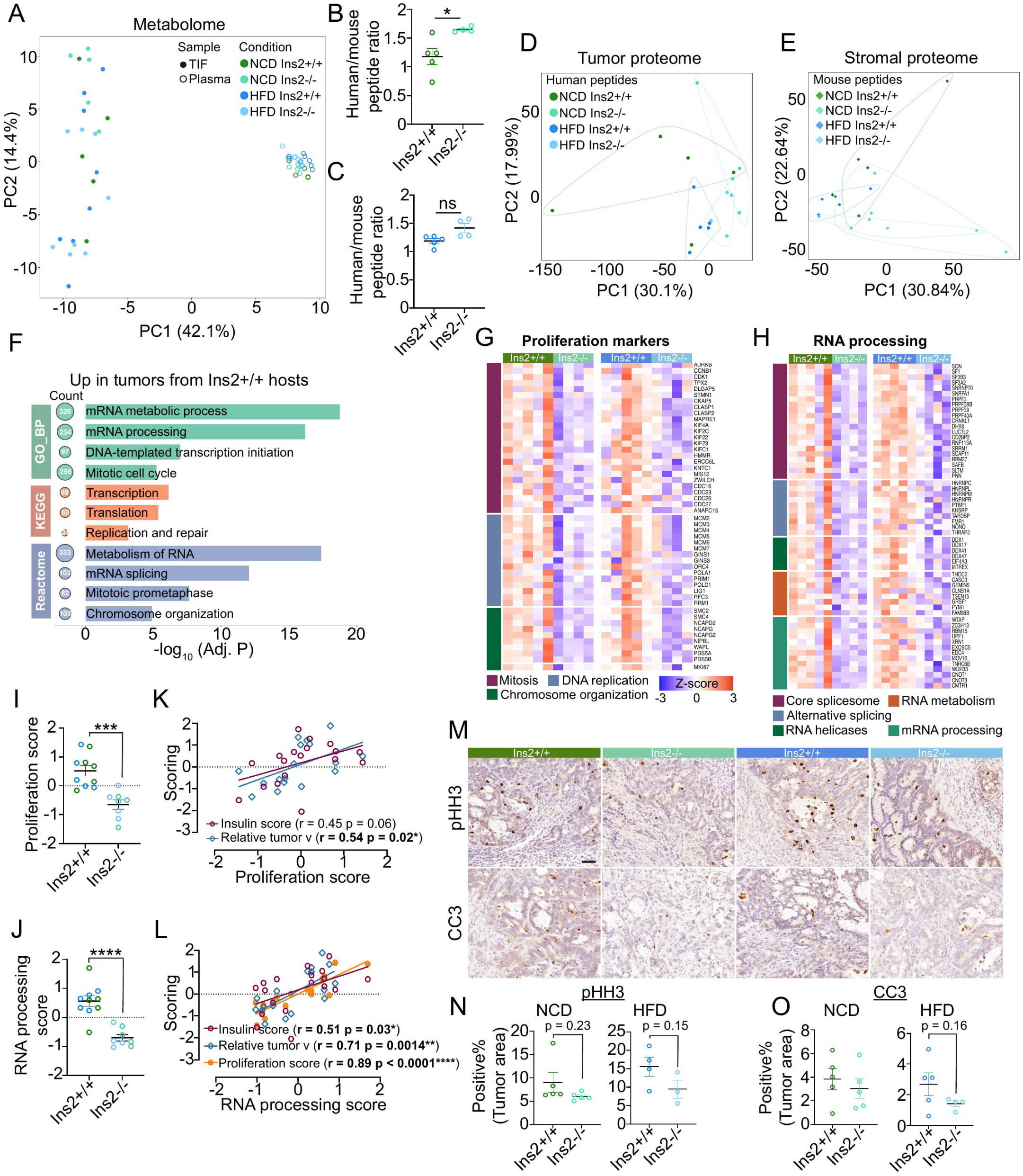
Reduced insulin is associated with reduced PDAC proliferation. (A) Principal component analysis (PCA) of polar metabolites measured in plasma or TIF samples from chow-fed and HFD-fed male cohorts. n = 5 in each NCD group; n = 8 in each HFD group. (B-C) The ratio of total human-specific peptides to total mouse-specific peptides in each tumor from the chow-fed (B) and HFD-fed (C) male mice. Values shown as mean ± SEM *p < 0.05 by two-tailed Student t-test. (D-E) PCA analysis of human/tumor-(D) and mouse/stroma-specific peptides. The top 500 most variable peptides were used in the analysis. (F) Over-representation analysis (ORA) of upregulated proteins in Fig. S6H. (G-H) Heatmap showing the relative abundance of proliferation marker proteins (G) and RNA-processing proteins (H). Z-score normalization was done within each diet condition. The protein panels were derived from the ORA analysis (F). (I-J) Proliferation (I) and RNA-processing (J) scores calculated by the mean of the protein z-scores of each sample (G-H). Values shown as mean ± SEM. ***p < 0.001, ****p < 0.0001 by two-tailed Student t-test. (K-L) Correlation of the proliferation score (K) and RNA-processing score (L) with the insulin score, relative endpoint tumor volume, or each other, pooling samples from both diet conditions together. r and p values were determined by Pearson correlation analysis. (M) Immunohistochemistry of phospho-histone 3 (pHH3) and cleaved-caspase 3 (CC3) for *Ins2^+/+^* and *Ins2^−/−^* tumor tissue. Scale bar = 50 μm. (N-O) Quantification of the % of tumor cells that are positive in pHH3 (N) or CC3 (O) in both diet conditions. Values shown as mean ± SEM.

Next, we focused on the changes in the proteome present in the tumor. For these data, we took advantage of the species difference between the immune/stromal and tumor components to selectively quantify the differences in each compartment in response to *Insulin* gene changes (Methods). This analysis yielded reliable quantitative data for 5,864 human-specific and 4,473 mouse-specific peptides across all samples (Extended data table S5). We found the total proportion of human-specific peptides in tumors from *Ins2^−/−^*hosts was higher under NCD, and tended to be higher under HFD as well (Fig. 6B-C), suggesting that reduced insulin was associated with a reduced host contribution to the tumor. To account for the variation in cellular composition across samples, species-specific peptides were normalized by the total human-to-mouse peptide ratio for each sample. PCA of the combined human-specific or mouse-specific proteomes showed partial separation of tumor proteomes by diet condition and genotype, while stromal-specific proteomes largely clustered together (Fig. 6D-E). Given the cell type heterogeneity of the stromal proteomes,^1^ we further examined whether differences could be seen in specific cell-type markers, but we did not see any clear differences (Fig. S6C-D). When we examined whether any of the mouse peptide changes correlated with mouse insulin score, we found 30 proteins that were consistently enriched with increasing insulin score and 5 that were decreased consistently across both diets (Fig. S6E-G). The presence of more mouse Chga and Krt19 in *Ins2^+/+^* tumors regardless of diet (Fig. S6F) suggested either random chance or a specific architecture of the tumor interface with the normal parenchyma led to the differences in the human-to-mouse peptide composition captured in the bulk piece of tumor. Alternatively, insulin could also have a concerted effect on the ratio of tumor proteins to stromal cells/matrix proteins that is shared across multiple cell types. Future single-cell-based analyses, particularly in an immune-competent background, would be necessary to fully address how insulin affects the tumor microenvironment.

Our further analyses focused on the insulin-dosage dependent differences in the cancer cell-specific proteomes. As expected from the overlap in the PCA plot, differential enrichment analysis between diets or genotypes did not identify any significantly differentially enriched peptides (Extended data Table S6). However, concordance analysis of the changes between tumors from *Ins2^+/+^* and *Ins2^−/−^* mice by diet (Fig. S6H) suggested a moderate, but significant, correlation of protein changes with *Ins2* gene dosage. Consistent with the cellular function of insulin to promote translation, *Ins2^+/+^-*derived tumors had approximately 3 times more proteins increased than decreased (Fig. S6H). Because these upregulated proteins were more consistently changed across samples, we investigated those proteins with consistent increases in tumors derived from *Ins2^+/+^* mice further. Ontology analysis suggested that these *Ins2^+/+^* up-regulated proteins were associated with proliferation, as well as RNA processing (Fig. 6F-H). As a group, we found that these proteins were generally lower in tumors grown in *Ins2^−/−^*mice (Fig.6I-J) and the relative Z-score for these protein groups across tumors correlated well with the relative tumor volume and insulin score, as well with each other (Fig 6K-L) This suggested that lowering insulin action/levels was associated with reduced PDAC cell proliferation. To validate this observation across additional samples, we performed immunohistochemistry against Serine 10 phosphorylation of histone 3 (pHH3), as well as the apoptosis marker, cleaved caspase-3 (CC3), to assess cell proliferation and survival. We found that the percentage of pHH3-positive tumor cells was ∼7% in NCD mice and 13% in HFD-fed mice supporting a role for HFD-induced hyperinsulinemia in promoting tumor cell proliferation. The percentage of pHH3^+^ tumor cells also tended to be higher in *Ins2^+/+^*compared to *Ins2^−/−^* tumors (Fig. 6M-N). In contrast, the CC3 positivity was less than 7% of tumor cells and varied widely across samples and genotypes but tended to be lower in the insulin reduced mice (Fig. 6M & 6O). Consistent with a role for insulin in cell proliferation in multiple cell types,^61–63^ we also found that mouse peptides for proliferation markers also tended to be higher in tumors from *Ins2^+/+^* compared to *Ins2^−/−^* mice (Fig. S6I). However, little to no pHH3^+^ or CC3^+^ cells were observed in stromal areas making validation difficult by IHC (Fig. 6M). In sum, increases in proteins associated with RNA-processing, likely due to increased cell proliferation, are associated with systemic increases in insulin-driven processes from before transplantation.

### Low expression of 45 proteins/transcripts associated with increased insulin score is associated with better patient survival

To identify proteins strongly associated with increased insulin activity, we next identified tumor proteins most strongly correlated with host insulin score across both diets (Fig. 7A). This analysis identified 45 or 4 tumor proteins that showed significant positive or negative correlations, respectively, with insulin score in both diet cohorts (Fig. 7A-B). The positively correlated proteins were involved in growth-associated functions, including mRNA processing, transcription, growth signaling, and mitochondrial support (Fig. 7B). Epithelial mucin and IGF signaling-related proteins showed negative correlation with insulin score (Fig. 7C), suggesting that tumors with reduced insulin may have PDAC cells where all the proteins needed for cell division/growth are less prominent. This would allow the epithelial-associated proteins like Muc1, Muc5B to be relatively enriched. Additionally, increased IGFBP3 may modulate the availability of IGF or Insulin in the tumor (Fig. 7C). To explore the clinical relevance of these insulin-associated proteins, we focused on the proteins associated with increased insulin action as they were more consistently changed in tumors from different genotypes (Fig. 7B). First, we used a single-cell RNA sequencing dataset comprising 24 human PDAC samples^64^ and used expression of the 45 positively correlated proteins (Fig. 7B) to group PDAC cells from each patient into high and low expression cell groups (Fig. 7D). By aggregating these high and low expressing cells for each patient into two data points (Pseudobulk), we could perform differential gene expression analysis of the high vs. low expressing cells across patients (n=24). This analysis confirmed that all 45 signature genes were higher in the signature-high group (Fig. 7E) an additional 362 and 297 genes were significantly upregulated or downregulated, respectively, in signature-high PDAC cells (Fig. 7E). Upregulated genes (not including the 45 insulin signature genes) were associated with cell proliferation, DNA replication, and RNA processing (Fig. 7F), whereas downregulated genes were associated with immune activation, apoptosis, and a catabolic state (Fig. 7G). Moreover, compared with signature-low PDAC cells, signature-high cells were predicted by PROGENy analysis^65^ to have higher PI3K, TGFβ, and MAPK signalling activity and lower activity of immune- and cell death-related pathways, including JAK-STAT and TRAIL signalling (Fig. 7H). Together, these analyses suggested that PDAC cells expressing higher transcript levels of insulin-associated proteins may be more proliferative and less immunogenic. Finally, we evaluated whether higher expression levels of the 45 genes correlated with high insulin activity were associated with differences in overall patient survival in the TCGA PAAD cohort.^66^ This analysis showed that patients whose tumors had lower expression of these 45 genes had significantly better overall survival than those with higher expression (Fig. 7I). Consistent with this relationship reflecting insulin’s role in promoting proliferation, the proliferative markers identified in our analysis (Fig. 6G) were also indicative of shorter survival times in the TCGA PAAD cohort (Fig. S7A). This suggests that readouts of relative insulin activity in the PDAC may be associated with patient outcomes.

**Figure 7.**
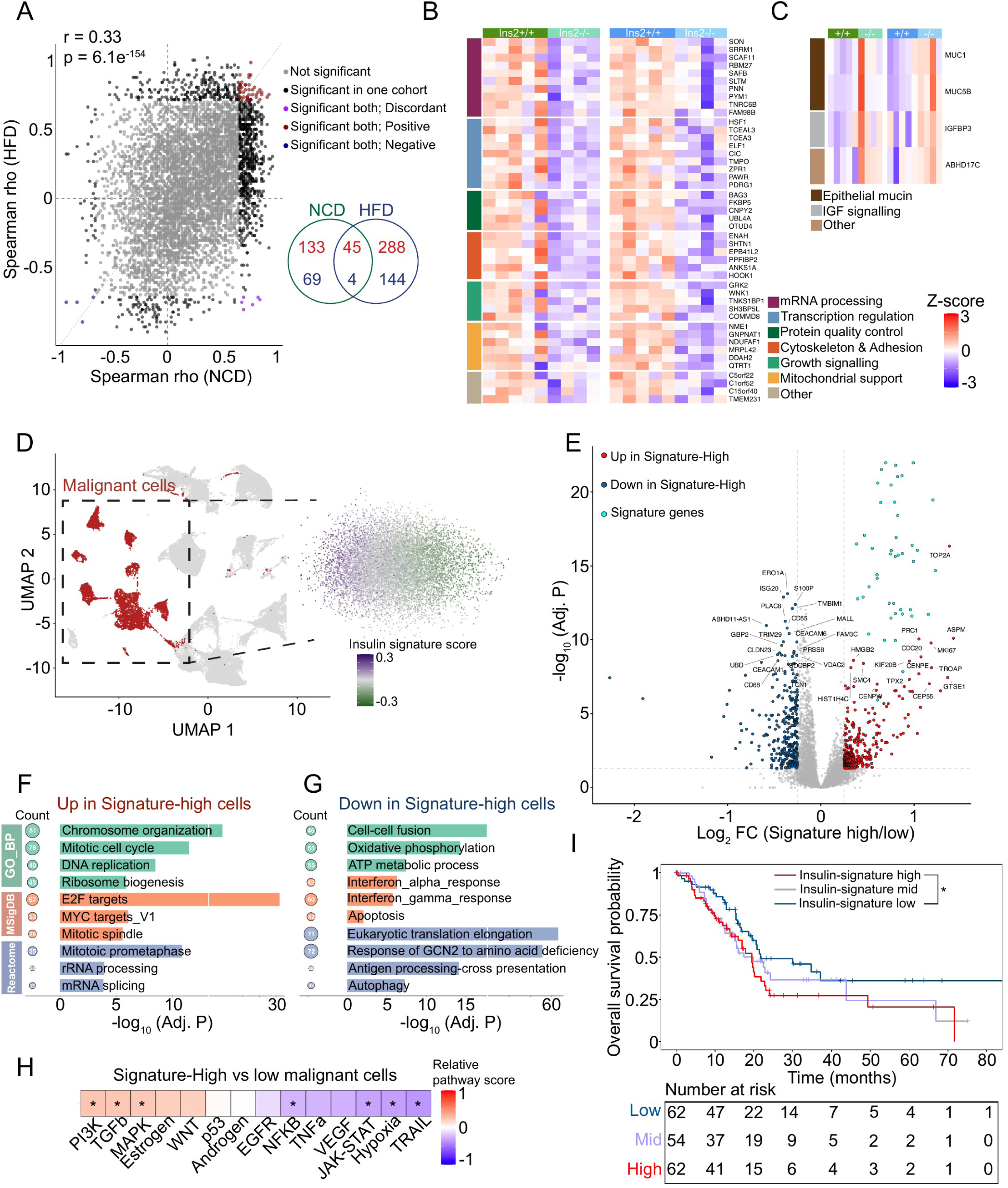
Proteins associated with increases in insulin activity in tumors are associated with poor survival outcomes in patients. (A) Concordance analysis of the Spearman correlation between each PDAC protein and the corresponding insulin score in both diet conditions. (B-C) Heatmaps that summarize the proteins showing significant positive (B) and negative (C) correlation with the host’s insulin score. (D) UMAP representations of the malignant cell population from scRNA sequencing analysis of 24 human PDAC tumors. An insulin signature score was assigned to each malignant cell within the same sample. (E) Volcano plots showing the differentially expressed genes between signature-high and signature-low malignant cell populations from pseudo-bulk analysis. (F-G) Over-representation analysis of the upregulated (F) and downregulated (G) genes in signature-high vs low malignant cells in (E). Signature genes were excluded from the over-representation analysis. (H) Pathway activity scores (PROGENy analysis) comparing signature-high to signature-low malignant cells in the pseudo-bulk analysis (E). Signature genes were excluded from the analysis. Positive scores (red) indicate that a pathway is predicted to be more active in signature-high cells; negative scores (blue) indicate that a pathway is predicted to be less active in signature-high cells. *p < 0.05 by PROGENy multivariate linear model. (I) Kaplan-Meier survival analysis of patient (TCGA PAAD) tumors with high and low mean expression of the 45 insulin signature genes. *p < 0.05 by log-rank test.

## Discussion

Our study provides evidence that changes in insulin levels are one important mechanistic link that could explain the spectrum of metabolic syndromes observed in PDAC patients. Specifically, long-term insulin-inducing metabolic syndromes, such as obesity and hyperinsulinemia, are associated with increased risk of PDAC,^3–5^ while PDAC diagnosis is often associated with signs of reduced insulin action, classified as type 3C diabetes.^5–7,9^ PDAC is also known to have high prevalence of cancer-associated cachexia,^67^ which is a complex sequelae of events associated with weight loss and muscle and fat catabolism in a manner somewhat reminiscent of Type 1 diabetes, which is caused by insulin-deficiency.^6,9^ Pre-clinically, there is also evidence that insulin is important for PDAC initiation and progression with dietary interventions that are known to either increase or decrease insulin, increasing or decreasing PDAC initiation and progression.^20,23–27^ While much less pre-clinical evidence exists that addresses the effects of PDAC on the pancreas, studies have suggested that pancreatic cancer worsens glucose homeostasis by increasing insulin resistance in either pancreatic beta cells^68^ or peripheral tissues, such as fat and muscle.^69–72^ In contrast to many of these studies, we examined the effect of PDAC on physiologic factors in more mature mice over a time period in which mice, particularly Bl6 male mice, naturally become slightly hyperinsulinemic.^52,54,55^ We found tumor growth limited the expected hyperinsulinemic effects caused by mouse background or HFD diet and actually improved glucose clearance, potentially reducing the overall demand for insulin from the islets. Additionally, in mice fed NCD, conditions where insulin levels are typically low, increased tumor growth was associated with decreased circulating insulin and catabolism of fat and muscle. These conditions likely further promoted tumor growth due the combined action of the high local insulin facilitating uptake of the catabolically liberated micromolecules from peripheral tissues and providing additional building blocks for proliferation when dietary nutrients were limiting. While we did not observe significant cachexia-like weight loss (>10% weight loss) and our mice did not become hyperglycemic before the experimental endpoint of 10 weeks, as often observed in patients,^49,67,73^, our study design and data provide more granular detail for these complex changes in mature tumor-bearing mice and establish a framework for further evaluating the paraneoplastic effects of pancreatic cancer on the normal pancreatic parenchyma and peripheral tissues.

The main purpose of this study was to test the hypothesis that reducing insulin would limit growth of PDAC in vivo. Unlike previous versions of our genetic insulin reduction models,^18,40^ in these experiments we only deleted *Ins2.* This was expected to result in small reductions in fasting plasma insulin. Indeed, we observed small, measurable differences in insulin that did not always meet significance threshold, likely due to the high degree of variability in single insulin measures. However, these small changes in insulin still had outsized effects on tumor growth, particularly in male mice which are prone to greater increases in insulin. Female mice generally maintain greater insulin sensitivity and may have greater compensation from the *Ins1* allele when *Ins2* is deleted.^13,18,48,53^ This suggests we may not have observed significant insulin-associated changes in females because they were able to maintain insulin action in ranges more similar to wild-type females during our study time frame. However, we did see that mature *Ins2^−/−^*females had small differences when compared to wild-type mice when challenged with HFD and in this context tumor growth was different by genotypes. Future studies testing whether other physiologically relevant female hyperinsulinemic models, such as ovariectomy-induced insulin resistance which mimics menopause,^74^ are needed to examine the role of insulin in pancreatic cancer growth under conditions that are more relevant to the female life stage in which PDAC is most often diagnosed. In sum, our data strongly suggest that very modest reductions in insulin function can reduce tumor growth, pointing out potential new therapeutic opportunities because many insulin modulating therapies and lifestyle changes currently exist that could be implemented to modestly reduce insulin levels and reduce the risk of early PDAC growth.

Circulating insulin levels are highly variable and subject to changes in energy expenditure, overall food intake, stress, and circadian rhythm.^57,75–78^ Unlike most related studies that inferred systemic insulin levels or function using a single insulin or glucose measurement,^21,27,39,79,80^ we developed a composite insulin score, which integrated longitudinal fasting insulin measurements with physiological features associated with overall insulin exposure. Because this score integrates several features, including those reflecting the cumulative effect of insulin’s anabolic functions, it may better represent an individual’s relative insulin exposure levels within the same experimental cohort. Indeed, this insulin score could represent differences in metabolic state that reflect how individuals of the same genotype respond to nutrient and environmental input. Interestingly, our study identified 45 tumor proteins that were highly associated with increased insulin score. In human PDAC patients, tumor cells showing higher transcript expression of these signature proteins showed higher activity of pro-tumor and insulin signalling effector pathways, and patients with higher expression of these 45 proteins had worse overall survival. Because the insulin score was calculated from pre-transplantation metrics, but was strongly correlated with the endpoint tumor volume, these data imply that an individual’s metabolic state may be a strong predictor of better tumor growth. Unfortunately, calculating a similar type of composite insulin score for patients with pancreatic cancer would not be feasible. Our longitudinal tracking of the insulin-related physiologic outputs demonstrated that these measurements changed as tumors grew in the host (as discussed above). Therefore, these differences across tumor cells from the same patient in the scRNAseq data and in the bulk samples used in the TCGA dataset, likely represent the propensity of a random tumor piece having high local areas of cell proliferation which in turn might be reflective of higher local insulin levels.

The human-specific tumor proteome indicates that reduced host insulin exposure is most consistently associated with downregulation of proliferation markers and RNA processing proteins, suggesting that the reduced tumor growth was primarily due to diminished PDAC cell proliferation. These data are consistent with the well-characterized role of insulin in cell proliferation.^30,32,62,81–83^ However the majority of these previous studies examined the acute effects of insulin on downstream insulin signaling effectors and cell proliferation or survival in 2D cell culture models,^81,82,84–90^ which have well-known dependencies on glycolysis.^91^ Organotypic or in *vivo* growth, however, is known to be less dependent on glucose.^80,91,92^ Therefore, our studies provide evidence that the physiologically relevant changes in insulin that are found in the pancreas under HFD or NCD conditions are important for promoting PDAC cell proliferation.

### Limitations

Although our mouse model provides a powerful platform to study the direct effects of insulin reduction in the context of human PDO growth, the *Rag1^−/−^* background lacks adaptive immunity. Thus, the effects we see occurred in the absence of functional B or T cells. However, insulin also has a key role in regulating wound healing through its effects on the immune cells and fibroblasts.^58,62,93^ As a chronic wound, the immune and stromal cellular components are also likely affected by insulin. As evidence of this, we observed differences in the ratio of human to mouse peptides in hosts with genetic reductions in insulin. Future studies that utilize spatial- or single cell-based approaches will be necessary to interrogate the influence modest changes in insulin have on the surrounding tumor microenvironment.

Systemic insulin homeostasis and sensitivity are well known sexually dimorphic traits.^46,48,53^ Both biological sexes were included and analyzed separately in our study with sex-matched PDO transplantation. Our physiological characterization revealed differences in the degree of metabolic changes observed in response to insulin reduction, reflecting sex differences in metabolism, as well as potential differences in compensation between sexes from the remaining *Ins1* alleles or other insulin-sensitive organs. However, since the male and female transplantations used different PDO lines, the difference between observations in males and females may also reflect, to some degree, tumor-intrinsic differences. Future studies using additional independent PDO lines from each sex are needed to elucidate whether heterogeneity exists in the response of tumors to reduced insulin, as well as in the heterogeneity of the tumors’ effects on the host physiology.

Finally, our metabolomic profiling focused on polar metabolites and did not include lipidomics. Since insulin is a central regulator of lipid synthesis, storage, and lipolysis in adipocytes and other cell types,^30,31^ altered systemic insulin exposure could affect lipid abundance and composition in circulation and TIF even when polar nutrient availability is not consistently genotype-dependent. This is particularly important because fat mass gains or losses were more sensitive readouts of overall insulin exposure. Future lipidomic profiling will be needed to determine whether insulin-mediated lipid remodelling contributes to PDAC growth suppression under reduced insulin conditions.

## Acknowledgments

The authors thank the Pancreas Center BC for PDOs. We thank Liam Hall, Shilpa Patil, Brian Daly, Anni Zhang, and Nan Chen for the training and discussion of mouse experiments. The project was supported by a MSFHR Scholar award and CIHR grant (#33524) to J.L.K. and by a Lustgarten Foundation Therapeutics Focused Research Program grant to J.D.J., D.F.S., J.L.K, and D.J.R. J.S.H.L was supported by a Four-Year Fellowship from the University of British Columbia and a Canada Graduate Research Scholarship (CIHR, #578083). D.F.S and D.J.R were supported by TFRI grant (#1078) and CCS grant (#706334). J.M.P and J.W were supported by a Canada Research Chair program grant (F18-01336). This work was also supported by grants to C.H.B from Génome Canada and Génome Québ́ec. The purchase of the Tims-TOf was supported by funding from the Canadian Foundation for Innovation grant (CFI-39858) to C.H.B. C.H.B is also grateful for support from the Segal McGill Chair in Molecular Oncology at McGill University (Montréal, Québec, Canada). C.H.B is also grateful for support from the Alvin Segal Family Foundation for the Segal Cancer Proteomics Centre at the Jewish General Hospital (Montréal, Québec, Canada), and for support from the Warren Y. Soper Charitable Trust for the Warren Y. Soper Clinical Proteomics Centre at the Jewish General Hospital (Montréal, Québec, Canada).

## Methods

### Mouse strains and breeding

University of British Columbia Animal Care Committee in accordance with Canadian Council for Animal Care guidelines approved all animal experiments. Mice were kept at the University of British Columbia Center for Disease Modelling (CDM). *Rag1^−/−^* mice were purchased from The Jackson Laboratory (JAX:002216). *Ins2^−/−^* mice have been previously described.^13,18,40^ *Rag1^−/−^* mice were bred with *Ins2^−/−^* mice to generate *Rag1^−/−^*; *Ins2^+/−^* breeders. Mating of these breeders created background-matched litters with the control (*Rag1^−/−^; Ins2^+/+^*) and experimental (*Rag1^−/−^; Ins2^−/−^*) genotypes. All mice used in each diet cohort, including the sham-transplanted mice, were generated in the same batch of mating, and the data were therefore normalized within each cohort. The number of mice measured across the cohort can vary depending on how many mice of the cohort were kept in the vivarium space where the necessary equipment was kept; movement between spaces was not allowed.

### Diet interventions and adult-stage paradigm

In the first *Ins2^+/+^* vs *Ins2^−/−^* cohort, the experimental litters were weaned (3 weeks) to a high-fat diet (HFD) with 60% fat (Research Diets D12492i; Research Diets). The mice received PDO transplantation at 10 weeks of age, with the surgical procedures outlined below. Tumors were collected and analyzed 10 weeks after transplantation. In the revised, adult paradigm HFD cohort, the experimental litters were weaned to a normal chow diet (NCD, Inotiv Teklad Global rodent diet 2918). The body composition of all mice was tracked weekly by a digital scale (body weight) or a body composition analyzer of fat and lean mass, EchoMRI-100 (EchoMRI-100; EchoMRI). The NCD was switched to an HFD once the mice stopped gaining lean mass. The mice received PDO transplantation 6 weeks after HFD initiation, with the surgical procedures outlined below. Tumor growth was monitored by the Visual Sonics Vevo2100 High-Resolution Ultrasound System (FUJIFILM VisualSonics Inc, VS-11945) weekly, as described by Goetze et al. 2018.^94^ Tumors were collected and analyzed 10 weeks after transplantation. Body weight, body composition, fasting plasma insulin, and fasting blood glucose were tracked throughout the entire experimental duration. Intraperitoneal insulin (ITT) and glucose (GTT) tolerance tests were performed at multiple time points before and after transplantation.

### Assessment of systemic insulin and glucose homeostasis

Fasting blood glucose and insulin were measured every 2-3 weeks. Mice were fasted for 4 hr before the blood collection for glucose/insulin measurement. The blood was collected from the saphenous vein. Glucose was measured with the Lifescan OneTouch Ultra Mini glucometer. To measure plasma insulin or C-peptide, ∼20uL of blood was collected into a heparinized microhematocrit capillary tube (Fisher Scientific, 22–362566), followed by 10 minutes of centrifugation at 10,000 rpm, 4 °C. The supernatant (plasma) was then collected and kept at −20 °C until the mouse insulin (ALPCO Diagnostics, 80-INSMSU-E10) or C-peptide (ALPCO Diagnostics, 80-CPTMS-E01) ELISA assay. Intraperitoneal GTT and ITT tests were performed as previously described.^13,40^. In brief, mice were fasted for 4 hr, followed by intraperitoneal glucose (GTT) or insulin (ITT, Eli Lilly, Cat. No. 0002-7510-01) injection at time zero. Blood glucose was measured before injection, and 15-, 30-, 60-, 90-, and 120 minutes after the injection. For GTT, 1.5 g/1 kg glucose was used for all conditions. For ITT, 0.5U insulin/kg was used for males under chow-feeding and females at all conditions. A higher dose (0.75U insulin/kg) was used for males under HFD-feeding. A composite insulin score was generated for each mouse by integrating longitudinal metabolic measurements to capture chronic systemic insulin exposure. Mean fasting insulin concentrations (HFD males: weeks 18, 20, 26; NCD males: weeks 17, 21, 26; HFD females: weeks 14, 16, 18; NCD females: weeks 17, 19) across at least two time points were log-transformed and within-cohort standardized (z-score), and combined with the pre-tumor z-scored mean of HOMA-IR (HFD & NCD males: weeks 18, 20, 26; HFD females: weeks 14, 16, 18; NCD females: weeks 17, 19), initial lean mass (HFD & NCD males: weeks 16-22; HFD females: weeks 15-19; NCD females: weeks 16-19) and the slope of fat mass accumulation over time (NCD males: weeks 16-22; NCD females: weeks 16-19). For the HFD cohorts, measurements collected after the diet switch, when fat mass accumulation had reached a relatively stable linear phase (HFD males: weeks 17-22; HFD females: weeks 14-19), were used to derive the fat mass slope component. The insulin score was calculated as: Insulin Score = 0.60 × Z(log10 Insulin) + 0.10 × Z(HOMA-IR) + 0.15 × Z(Lean Mass) + 0.15 × Z(Fat Mass Slope). The weighting scheme emphasizes circulating insulin while incorporating insulin resistance and body composition parameters that are known to reflect cumulative insulin exposure. Higher insulin scores are expected to reflect greater chronic insulin exposure. This score was used as a continuous variable in correlation analyses to evaluate associations between insulin exposure and tumor growth or proteomic changes.

### Culture of Patient-derived PDAC organoids (PDOs)

PDOs were generated at the Pancreas Center BC.^35^ The PDO culture protocol was modified from the Tuveson protocol.^95^ PDOs were resuspended in 100% growth factor reduced Matrigel (Corning, 354230) and plated 50 *μ*L per well in a 24-well plate as domes. Domes were overlaid with the completed growth media. The basal growth media (control condition) was formulated as: 50% Human plasma-like media (Gibco, Cat. No. A4899102) supplemented with 1X GlutaMAX (ThermoFisher Scientific, 35050061), 1x HEPES (ThermoFisher Scientific, 15630080), 100 *μ*g/mL primocin (Invivogen, ant-pm-1), 1X B-27 supplement minus insulin (Thermo FisherScientific, A1895601), 1.25mM N-Acetyl-L-cysteine (Sigma-Aldrich, A9165), 10nM hGastrin-I (Sigma-Aldrich, G9020), 100 ng/mL Recombinant hFGF-10 (Peprotech, 100 − 26), 0.5 *μ*M A 83 − 01 (Tocris, 2939),10*μ*M Y-27632 (Tocris, 1254), 10 *μ*M Nicotinamide (Sigma-Aldrich, N0636), recombinant hEGF (Peprotech, AF-100-15), and 50% Wnt-3a/R-Spondin1/Noggin conditioned medium (ATCC, 3276).

### Orthotopic transplantation of patient-derived tumor organoid (PDO)

The surgical procedure was adapted from the protocol described by previous literature.^96^ Briefly, the mice were anesthetized with 4-2% isoflurane with a partial pressure of 400 cc O_2_. Fur was shaved around the skin where the pancreas is located. The mice were provided eye lubricant (AVP, 2122859), 5 mg/kg meloxicam (MODERN VET, 3820225Y), 0.05 mg/kg buprenorphine (Ceva, 124918), 8 mg/kg bupivacaine hydrochloride (HENRY SCHEIN, 1130135), Enrofloxacin (ELANCO, 2580164Y), and 0.5-1 mL Lactated Ringer’s Solution (BAXTER, 1672237). A left flank incision was made, followed by a gentle externalization of the spleen and the pancreas tail. Approximately 5,000 (PCBC5, female) or 10,000 (PCBC8, male) PDO cells suspended in 20 μL undiluted Matrigel solution were injected into the tail of the pancreas with a 29G insulin syringe. Sham-transplanted animals received orthotopic injection of empty Matrigel solution. After the injection, the spleen and pancreas were re-placed, and the peritoneum and skin were closed with Monocyl sutures (J&J, 1651307). After the surgery, the mice were given buprenorphine for 3 continuous days (twice a day, 12 hours gap between each injection) and monitored until the incision was healed and body weight was stable.

### Necropsy and sample collections

Mice were weighed immediately after euthanasia. Endpoint plasma was collected through cardiac puncture. For assessment of normal tissue weights, the quadriceps and gastrocnemius muscles from the left hindlimb, subcutaneous (left), perigonadal (left), retroperitoneal (right), and brown (shoulder) adipose tissues, liver, and heart were dissected and weighed. The tumor was dissected, measured, and weighed, and then the tumor interstitial fluid (TIF) isolation procedure adapted from Sullivan et al. 2019^97^ was performed. Briefly, a random piece of tissue was collected from each tumor and placed on a 1.5mL microcentrifuge tube covered by a piece of nylon filter (Spectrum Labs, 148134). The construct was centrifuged at 4°C for 10 min at 106 × g. After centrifugation, the fluid at the bottom of the conical tube was collected, snap-frozen in liquid nitrogen, and kept at −80 °C until metabolomics or insulin/glucose analysis. PanIF was isolated from the normal adjacent portion of tumor-bearing pancreas (tail region) or the pancreatic tail from sham-transplanted mice. One random piece of tumor tissue was collected and snap-frozen in liquid nitrogen and kept at −80 °C until proteomics analysis.

### Histopathological, morphological, and immunohistochemical analyses

Tissues were fixed in 4% paraformaldehyde for 24 hrs at 4 °C, followed by processing (Leica, ASP300S), paraffin embedding (Leica, ArcadiaH), and sectioning at 7 μM. The sections were stained with hematoxylin and eosin (H&E) or immunohistochemistry (IHC) as described.^10,40^ Primary antibodies used in IHC were rabbit anti-Phospho-Histone H3 (Ser10) (Cell Signalling, 9701, 1:500) and rabbit anti-Cleaved-caspase 3 (cleaved Asp175) (ThermoFisher, PA5-114687, 1:200), followed by biotin-conjugated donkey anti-rabbit (Jackson Laboratory, 711-065-152, 1:500) secondary antibody. The stained slides were scanned with a 20x objective using a 3DHISTECH Panoramic MIDI (Quorum Technologies, Guelph, Canada) slide scanner. Percent positive IHC stains were quantified using QuPath 0.5.1.

### Protein extraction for Mass spectrometry-based proteomics

Frozen tumor specimens underwent tissue lysis by bead beating in CK-14 tubes with 200 μL of tissue extraction buffer containing 5% SDS, 100 mM TRIS pH 8.5, and 10 mM TCEP in a Precellys CryoLys system (Bertin) with 3 cycles of shaking for 20 seconds at 6500 rpm. Lysates were further extracted by heating at 95°C for 10 minutes with rotational agitation at 1800 rpm using an Eppendorf ThermoMixer C. Lysates were then clarified by centrifugation at 21,000 x g for 1 minute, and the supernatant was transferred to new microcentrifuge tubes. Protein concentrations were determined using a reducing agent-compatible bicinchoninic acid assay kit (Thermo Fisher Scientific, Cat. No. 23250). The remaining sample was alkylated with 40 mM iodoacetamide for 20 minutes in the dark. An equivalent of 20 µg of protein was proteolytically digested with sequencing-grade trypsin (Promega, Cat. No. V5111) at 37 °C overnight (16 hours) using S-TRAP microcartridges according to the vendor’s protocol (Protifi LLC).

### LC-MS/MS proteomics analysis

For global proteomics, an equivalent of 200 nanograms per sample was loaded onto Evotip Pure tips (Evosep) according to the manufacturer’s protocol and analyzed using an Evosep One LC and a Bruker timsTOF HT mass spectrometer operated in DIA-PASEF mode using an empirically optimized isolation scheme using the py_diAID tool (https://github.com/MannLabs/pydiaid) using 12 PASEF ramps split with 24 MS/MS windows, and had an estimated cycle time of 1.2 seconds. Precursor isolation was between 100 – 1700 m/z and an ion mobility range of 0.7 – 1.4 1/k0, with ramp and accumulation times set to 100 ms. Collision energy was scaled based on precursor ion mobility from 20 – 59 eV between 0.6 and 1.6 1/k0, respectively. DIA-PASEF data was analyzed using DIA-NN (2.3.1) using an in silico predicted spectral library generated by DIA-NN, based on the canonical human and mouse reference proteomes (UP000005640, UP000000589) and the following application settings: MBR enabled, protein inference selected, proteotypicity based on species, cross run normalization based on retention time. Protein group-level quantitation (report.gg_matrix.tsv) was used for downstream statistical analysis.

### Tumour interstitial fluid (TIF) and plasma analysis using liquid chromatography-mass spectrometry metabolomics

Previously frozen TIF and plasma samples were thawed on ice then centrifuged (4°C) for 5 min at 21,300xg. One-five µL of TIF/plasma samples were then added to 250 µL of ice-cold extraction buffer. The extraction buffer consisted of 80% methanol and 20% water (v/v) supplemented with a stable isotope labeled internal amino acid standard mix (w/o L-Trp) (Cambridge Isotope Labs, MSK-A2), stable isotope labeled tryptophan (CDN Isotopes, D-1522), stable isotope labeled glucose (Cambridge Isotope Labs, CLM-481), and stable isotope labeled lactate (Cambridge Isotope Labs, CLM-1579), all at 1 nmol per sample. Samples were shaken for 15 minutes (1500 rpm), centrifuged (4°C) for 15 min at 21,300xg, and the supernatant (∼240 µL) transferred to a new tube. Samples were dried to completion overnight using a SPD120 SpeedVac (Thermo Fisher). Dried metabolites were reconstituted in 25 µL of water, shaken for 10 minutes (1500 rpm), centrifuged (4 °C) for 5 min at 21,300xg, and the supernatant was transferred to LCMS vials for analysis. Vials were loaded into a temperature-controlled (6°C) autosampler of a Vanquish LC (Thermo Scientific). The autosampler was coupled to a ZIC-pHILIC LC column, with a volume oven temperature set to 25°C, maintained by forced air and an integrated column heater. The mobile phase solvents were: (A) 10 mM ammonium carbonate in HPLC-grade water, pH 9.0; (B) 100% acetonitrile. The column was pre-equilibrated using a flow rate of 100 μL/min and 80%(B)/20%(A). Following injection of 2 μL of sample, the following gradient for elution at 100 μL/min based on % of (A) was used: 20-80%(A) (0–30 min), 80-80%(A) (30–40 min), and 80–20%(A) (40–40.5 min); the LC column was re-equilibrated using high mobile phase (B), 80-80%(B) from 40.5 to 52 min before subsequent injections. Liquid chromatography was coupled to an Exploris 240 mass spectrometer (Thermo Scientific) operating in heated electrospray ionization (HESI) mode. HESI source parameters were as follows: spray voltage 3.4 kV (positive mode) and 2.0 kV (negative mode), static spray voltage; sheath gas 25; auxiliary gas 5; sweep gas 0.5; ion transfer tube temperature 320 °C; vaporizer temperature 75 °C. Global instrument parameters included an expected peak width of 20 s, mild trapping, and a default charge state of 1. Full MS scans (positive and negative modes) were acquired in profile mode over a scan range of 67–1000 m/z at 120,000 resolutions, with RF lens set to 70%, AGC target of 300%, and maximum injection time set to automatic. ddMS2 for positive mode were collected in centroid mode at an Orbitrap resolution of 30,000, isolation window of 1.5 m/z, an AGC target set to standard, a maximum injection time set to automatic, and a normalized collision energy set to 10%, 30%, and 80%. ddMS2 for negative mode were collected in centroid mode at an Orbitrap resolution of 30,000, isolation window of 2 m/z, an AGC target set to standard, a maximum injection time set to automatic, and a normalized collision energy set to 30%. Raw LCMS data were processed using TraceFinder (Thermo Fisher Scientific). Metabolites were annotated based on accurate mass and retention time matching to an annotated library generated from the Mass Spectrometry Metabolite Library of Standards and additional authentic standards obtained from Sigma. Quality control (QC) filtering of metabolite peaks was performed using minimum signal intensity and blank contamination thresholds. Signal intensity thresholds were determined from the mean peak intensity across samples in the dataset. Peaks were excluded if they fell below the minimum intensity threshold of ≤2.5 × 10⁵ AU or if the signal detected in blank controls exceeded the specified fraction of ≥20% of the sample signal average.

### Bioinformatic analysis

Human pancreatic ductal adenocarcinoma (PAAD) dataset CRA001160 was obtained from the TISCH portal. Malignant cells were annotated according to the TISCH malignancy annotation. The insulin-associated transcriptional program is defined by the 45 proteins positively associated with insulin exposure in our tumor proteomics dataset. For each cell, an insulin program score was calculated using a control-matched module scoring approach,^98^ where the mean expression of signature genes was corrected by the mean expression of control genes sampled from matched expression bins. Signature states (signature-high, top30%; signature-low, bottom 30%) were assigned among malignant cells in a patient-specific manner. Pseudobulk differential expression (DE) analysis was performed using edgeR with a paired patient-blocked design (∼ Patient + State) with FDR correction. Over-representation analysis and pathway activity of differentially expressed genes (signature-high vs low) were performed using clusterProfiler^99^ and PROGENy^65^ R packages and custom scripts. Overall survival analysis was performed in R using the survival and survminer packages and pancreatic ductal adenocarcinoma (PAAD) data from The Cancer Genome Atlas (TCGA), accessed through the UCSC Xena platform (n = 178). In Kaplan–Meier analysis, tumors were stratified according to the distribution of insulin signature scores into bottom 35%, middle 30%, and top 35% groups. Survival differences were evaluated between the bottom 35% and top 35% groups, and significance was assessed using a two-sided log-rank test comparing the bottom and top 35% groups.

### Statistical analysis

Statistical analyses were conducted with GraphPad Prism 10. Results are shown as mean ± SEM unless otherwise indicated. The Shapiro-Wilk and F tests were used to assess data normality and variances. The two-tailed Student t-test was used for normally distributed data, whereas the Mann-Whitney test was performed for non-normally distributed data when comparing the mean between two groups. The Welch t-test was used when two normally distributed populations have unequal variances. The Paired t-test was used when comparing two groups within matched samples. The Pearson correlation test was run to determine the correlation coefficient (r) and p values for linear correlation between two groups. Spearman correlation test (rho and p values) was run to determine monotonic relationships that may not be linear. Comparisons of multiple groups and/or conditions were done by one-way or two-way analysis of variance (ANOVA) when appropriate. The linear Models for microarray (LIMMA) R package^100^ was used for differential expression analysis for proteomics data. In figures, asterisks denote statistical significance (*p < 0.05, **p < 0.01, ***p < 0.001, and ****p < 0.0001).

## Declaration of generative AI and AI-assisted technologies

During the preparation of this work, the author(s) used ChatGPT for literature search (Deep research tool), coding assistance and debugging. The author(s) reviewed and edited the output as needed and takes full responsibility for the content of the published article.

---

**Figure S1.**
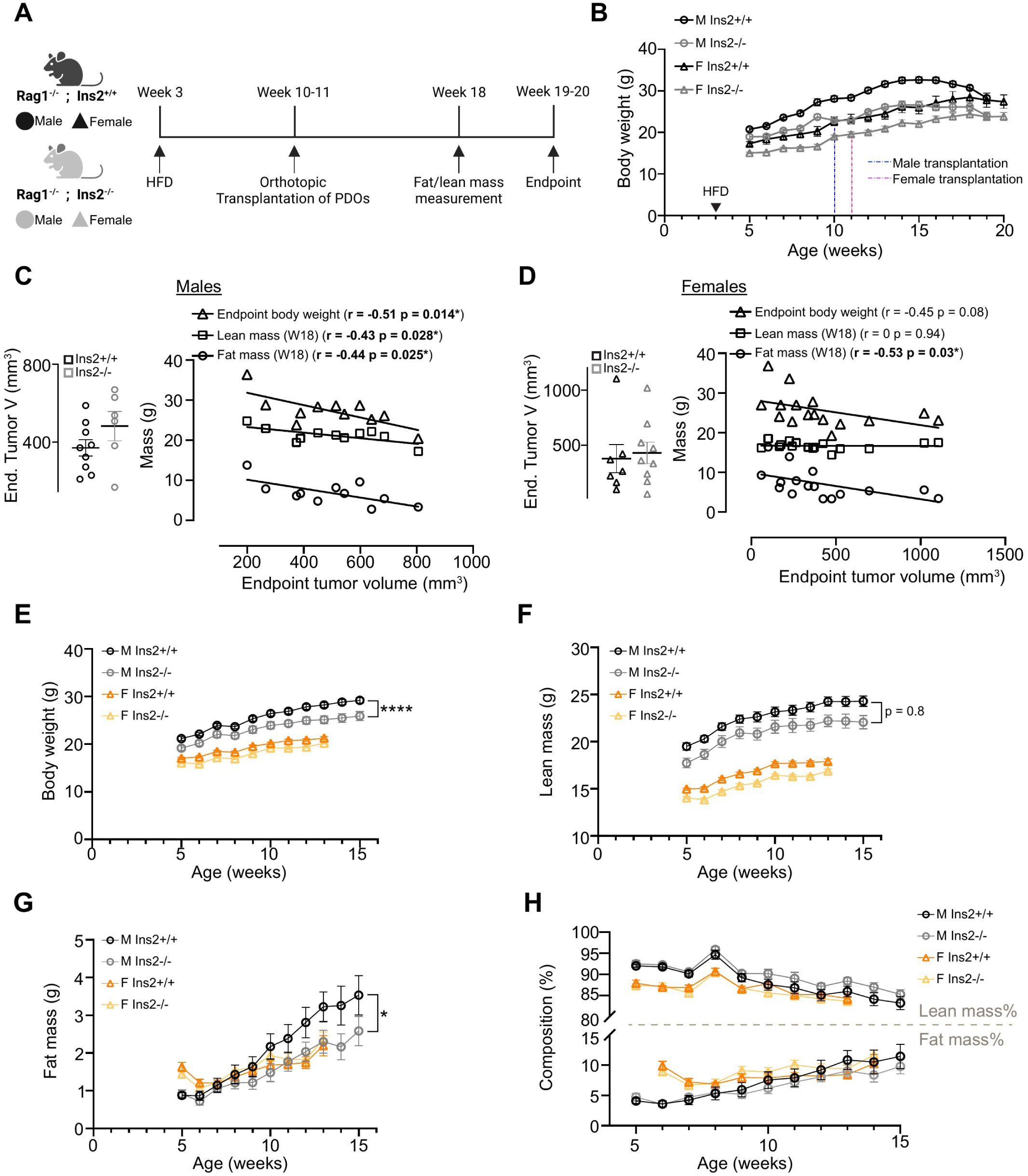
Genetic reduction in *insulin* affects body growth and tumor growth during this life stage confounds the influence of insulin, related to Figure 1. (A) Schematic describing the first experimental design to study orthotopic tumor progression in *Ins2^+/+^*and *Ins2^−/−^* mice under HFD-feeding. n = 5-7 in each group. (B) Body weight of experimental mice measured over time. Values at each time point are shown as mean ± SEM. (C) Left panel: Calliper-measured endpoint tumor volume in male mice. Values shown as mean ± SEM. Right panel: Correlation of endpoint tumor volume with endpoint body weight, week 18 lean mass, and week 18 fat mass in all male mice (correlations values shown in legend). Correlation by specific genotype: Body weight (*Ins2^+/+^*: r = −0.71, p = 0.03; *Ins2^−/−^*: r = −0.57, p = 0.42); Lean mass (*Ins2^+/+^*: r = −0.79, p = 0.03; *Ins2^−/−^*: r = −0.51, p = 0.45); Fat mass (*Ins2^+/+^*: r = −0.64, p = 0.12; *Ins2^−/−^*: r = −0.91, p = 0.09). r and p values were determined by Pearson correlation analysis. (D) Left panel: Calliper-measured endpoint tumor volume in female mice. Values shown as mean ± SEM. Right panel: Correlation of endpoint tumor volume with endpoint body weight, week 18 lean mass, and week 18 fat mass in all female mice (correlation values shown in legend). Correlation by specific genotype: Body weight (*Ins2^+/+^*: r = −0.55, p = 0.20; *Ins2^−/−^*: r = −0.32, p = 0.39); Lean mass (*Ins2^+/+^*: r = −0.10, p = 0.87; *Ins2^−/−^*: r = 0.13, p = 0.72); Fat mass (*Ins2^+/+^*: r = −0.62, p = 0.13; *Ins2^−/−^*: r = −0.43, p = 0.23). r and p values were determined by Pearson correlation analysis. (E-G) Body weight (E), lean mass (F), and fat mass (G) of chow-fed *Ins2^+/+^* or *Ins2^−/−^* male and female mice over time. (H) The percentage of lean mass and fat mass with respect to the total body weight of *Ins2^+/+^* or *Ins2^−/−^* male and female mice over time.

**Figure S2.**
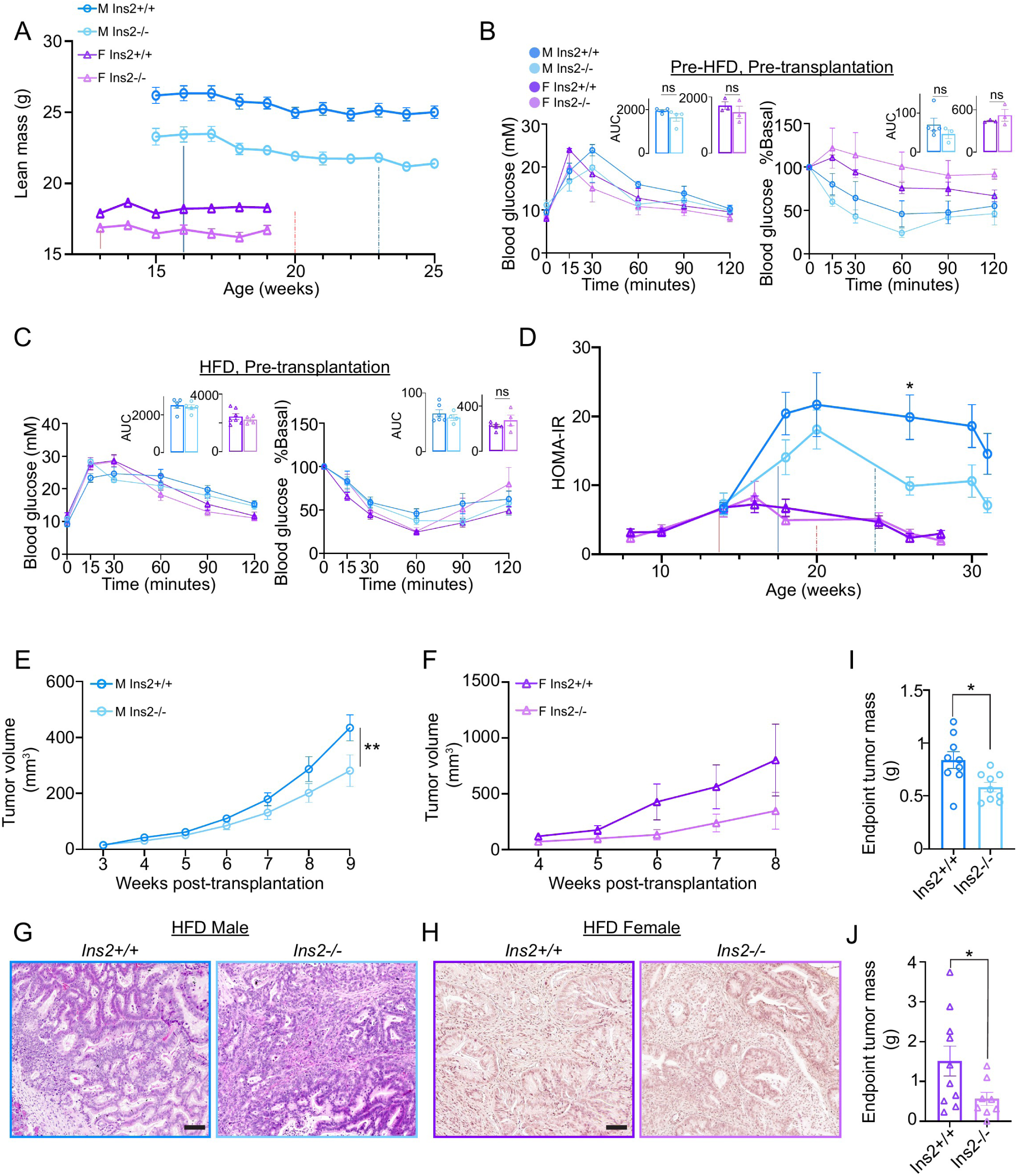
Effects of *Ins2* loss on systemic metabolism and PDAC growth in HFD-fed mice, related to Figures 2 and 3. (A) Lean mass of HFD-fed mice measured over time before and/or shortly after transplantation. Values at each time point are shown as mean ± SEM. (B-C) Glucose measures during an intraperitoneal glucose tolerance test (GTT) (left panel) or insulin tolerance tests (ITT)(right panel) performed with HFD-fed male and female mice at the following time points: Pre-HFD, pre-transplantation (B, week 16 for males and week 12 for females) and post-diet switch and pre-transplantation (C, week 21 for males and week 19 for females). Area under the curve (AUC) values shown as mean ± SEM. (D) HOMA-IR of HFD-fed mice measured over time. Values at each time point are shown as mean ± SEM. *p < 0.05 by one-way ANOVA. (E-F) Tumor volume over time of male (E, n = 6-7) and female (F, n = 5-8). (G-H) Representative H&E images of tumors from HFD-fed male (G) and female (H) mice of each genotype. Scale bar = 100 μm. (I-J) Tumor mass measured at the endpoint dissection of HFD-fed male (I) and female (J) mice across genotypes. Values shown as mean ± SEM *p < 0.05 by Welch t-test.

**Figure S3.**
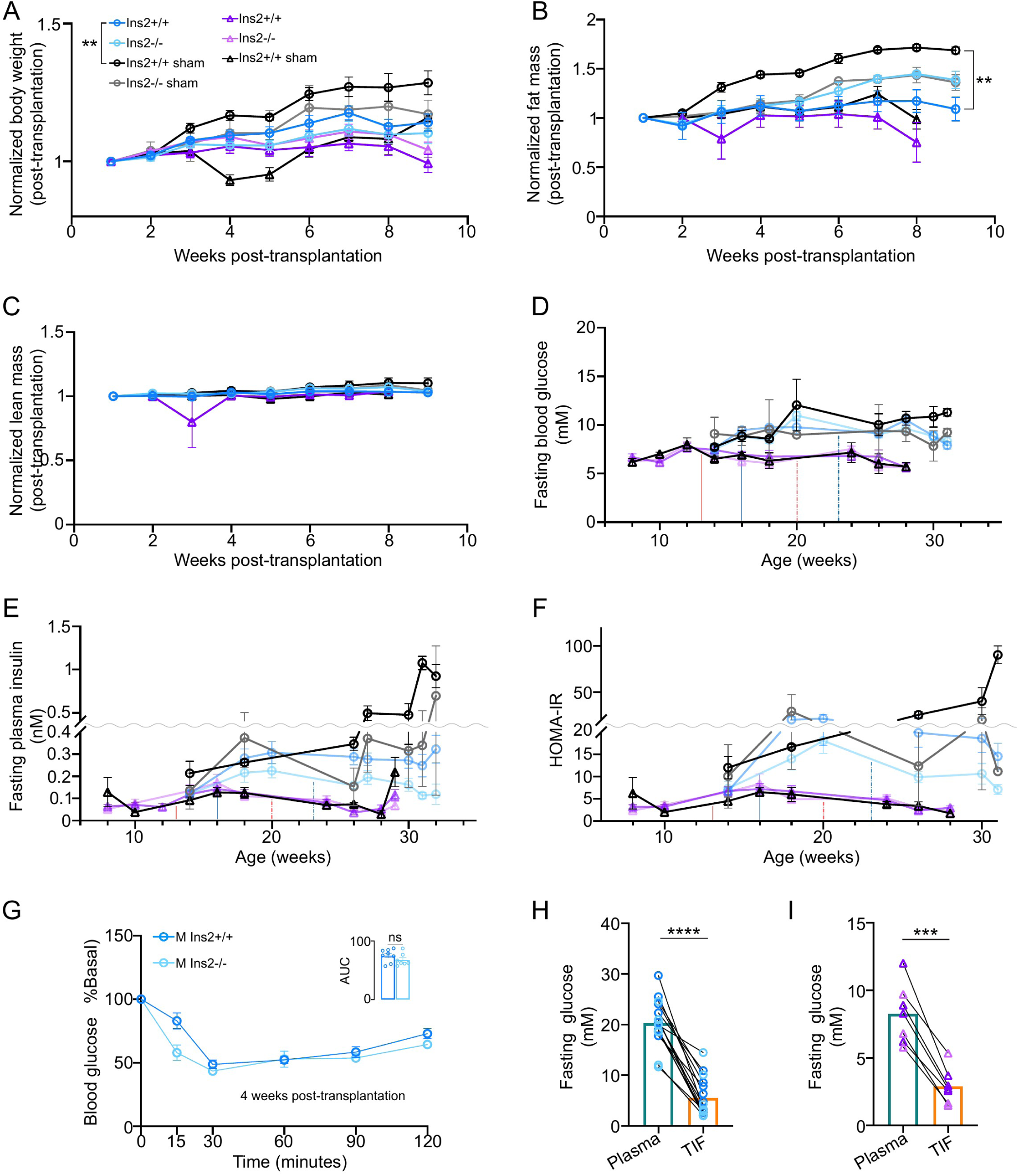
Presence of a tumor in the pancreas affects the expected HFD-induced effect on host metabolism, related to Figure 3. (A) Post-transplantation body weight of HFD-fed tumor- and sham-transplanted mice normalized to the body weight 1-week post-transplantation ((n = 10, *Ins2^+/+^* tumor male; n = 9, *Ins2^−/−^* tumor male; n = 10, *Ins2^+/+^* tumor female; n = 6, *Ins2^−/−^* tumor female; n = 3, *Ins2^+/+^*sham male; n = 2, *Ins2^−/−^* sham male; n = 3, *Ins2^+/+^*sham female). *Ins2^−/−^* sham was not evaluated in females. Values at each time point are shown as mean ± SEM. **p < 0.01 by mixed effects test of growth rate. (B) Post-transplantation fat mass of HFD-fed tumor- and sham-transplanted mice normalized to the fat mass 1-week post-transplantation ((n = 3, *Ins2^+/+^* tumor male; n = 2, *Ins2^−/−^* tumor male; n = 5, *Ins2^+/+^* tumor female; n = 3, *Ins2^+/+^* sham male; n = 2, *Ins2^−/−^*sham male; n = 3, *Ins2^+/+^* sham female). *Ins2^−/−^*tumor- or sham-transplanted females were not evaluated. Values at each time point are shown as mean ± SEM. **p < 0.01 by mixed effects test of growth rate. (C) Post-transplantation lean mass of HFD-fed tumor- and sham-transplanted mice normalized to the lean mass 1-week post-transplantation ((n = 3, *Ins2^+/+^* tumor male; n = 2, *Ins2^−/−^* tumor male; n = 5, *Ins2^+/+^* tumor female; n = 3, *Ins2^+/+^* sham male; n = 2, *Ins2^−/−^* sham male; n = 3, *Ins2^+/+^*sham female). *Ins2^−/−^* tumor- or sham-transplanted females was not evaluated. Values at each time point are shown as mean ± SEM. (D-E) Fasting blood glucose (D) and HOMA-IR (E) of HFD-fed sham-transplanted mice over time. Values at each time point are shown as mean ± SEM. Levels from tumor-transplanted mice (Fig. 2E and S2D) are plotted for comparison. Sample size information is consistent with Fig.S3A. (F) Intraperitoneal insulin tolerance (IPITT) tests performed with HFD-fed tumor-transplanted male mice 4 weeks post-transplantation (week 27). Area under the curve (AUC) values shown as mean ± SEM. (G) Fasting insulin of HFD-fed sham-transplanted mice over time. Values at each time point are shown as mean ± SEM. Levels from tumor-transplanted mice (Fig. 2D) are plotted for comparison. Sample size information is consistent with Fig.S3A. (H-I) Glucose concentrations in TIF and the matched plasma samples at fasting from HFD-fed male (H)(measured by LC-MS) and female (I) (measured by colorimetric assay) mice.

**Figure S4.**
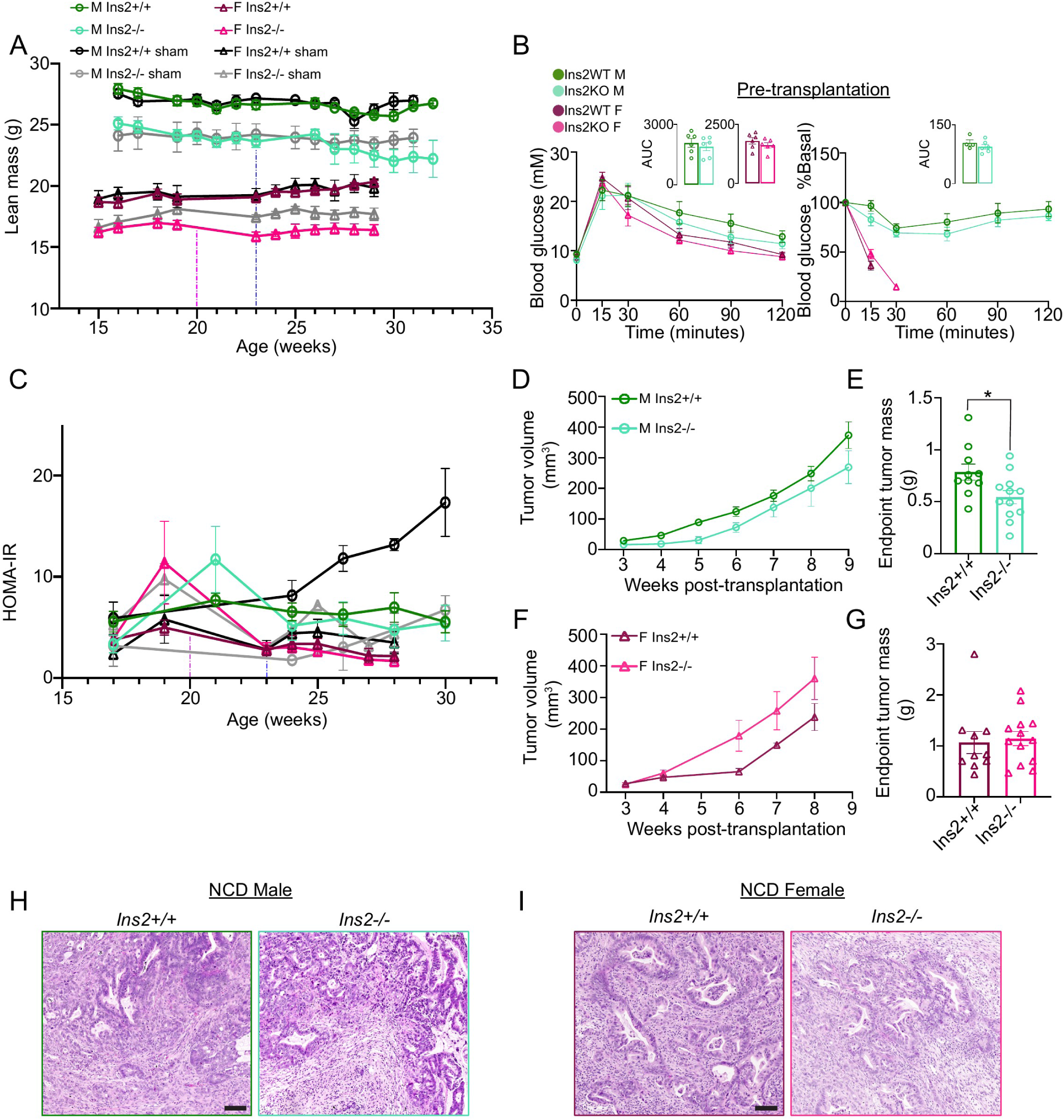
Genetic reduction of *insulin* affects PDAC growth in chow-fed mice, related to Figure 4. (A) Lean mass of chow-fed mice measured over time before and/ after transplantation. Values of tumor-transplanted mice were obtained from 3 animals (n=3 for each group of tumor-transplanted mice). Values at each time point are shown as mean ± SEM. (B) Intraperitoneal glucose (IPGTT) and insulin tolerance (IPITT) tests performed with chow-fed experimental male and female mice before transplantation (week 21 for males and week 19 for females. The female ITT experiment was terminated due to hypoglycemia). (C) HOMA-IR of chow-fed mice over time. Values at each time point are shown as mean ± SEM. (D-E) Tumor volume over time (n = 7-8) measured by ultrasound imaging (D, values at each time point are shown as mean ± SEM) and the calliper-measured tumor volume (E, values shown as mean ± SEM. *p < 0.05 by two-tailed Student t-test.) at the endpoint dissection of chow-fed male mice. (F-G) Tumor volume over time of female mice (n = 7-10) measured by ultrasound imaging (F) and the calliper-measured tumor volume (G) at the endpoint dissection of chow-fed female mice. (H-I) Representative H&E images of tumors from chow-fed male (H) and female (I) mice. Scale bar = 100 μm.

**Figure S5.**
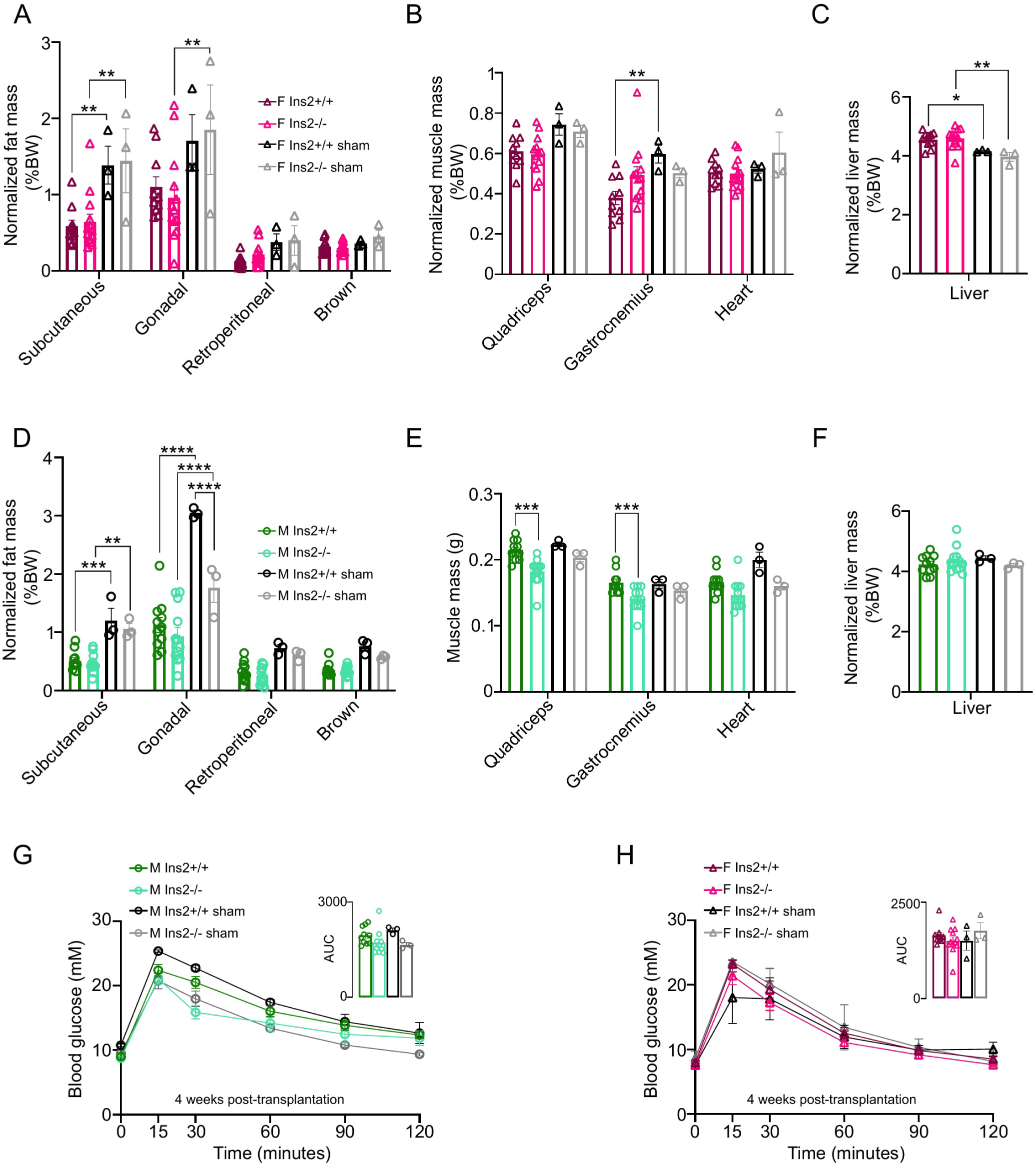
PDAC growth in NCD-fed mice is associated with reduced fat and lean mass, related to Figures 4 and 5. (A-C) The mass of adipose depots (A), muscle depots (B), and liver (C) of chow-fed sham- and tumor-transplanted females normalized by the endpoint body weight. Values shown as mean ± SEM. *p < 0.05, **p < 0.01 by two-way ANOVA. (D-F) The normalized mass of adipose depots (D), absolute mass of muscle depots (E), and normalized mass of liver (F) of chow-fed sham- and tumor-transplanted males.Values shown as mean ± SEM. **p < 0.01,***p < 0.001, ****p < 0.0001 by two-way ANOVA. (G-H) Intraperitoneal glucose (IPGTT) tests performed with sham- and tumor-transplanted male (G, week 27) and female (H, week 24) mice 4 weeks after transplantation. Area under the curve (AUC) values shown as mean ± SEM.

**Figure S6.**
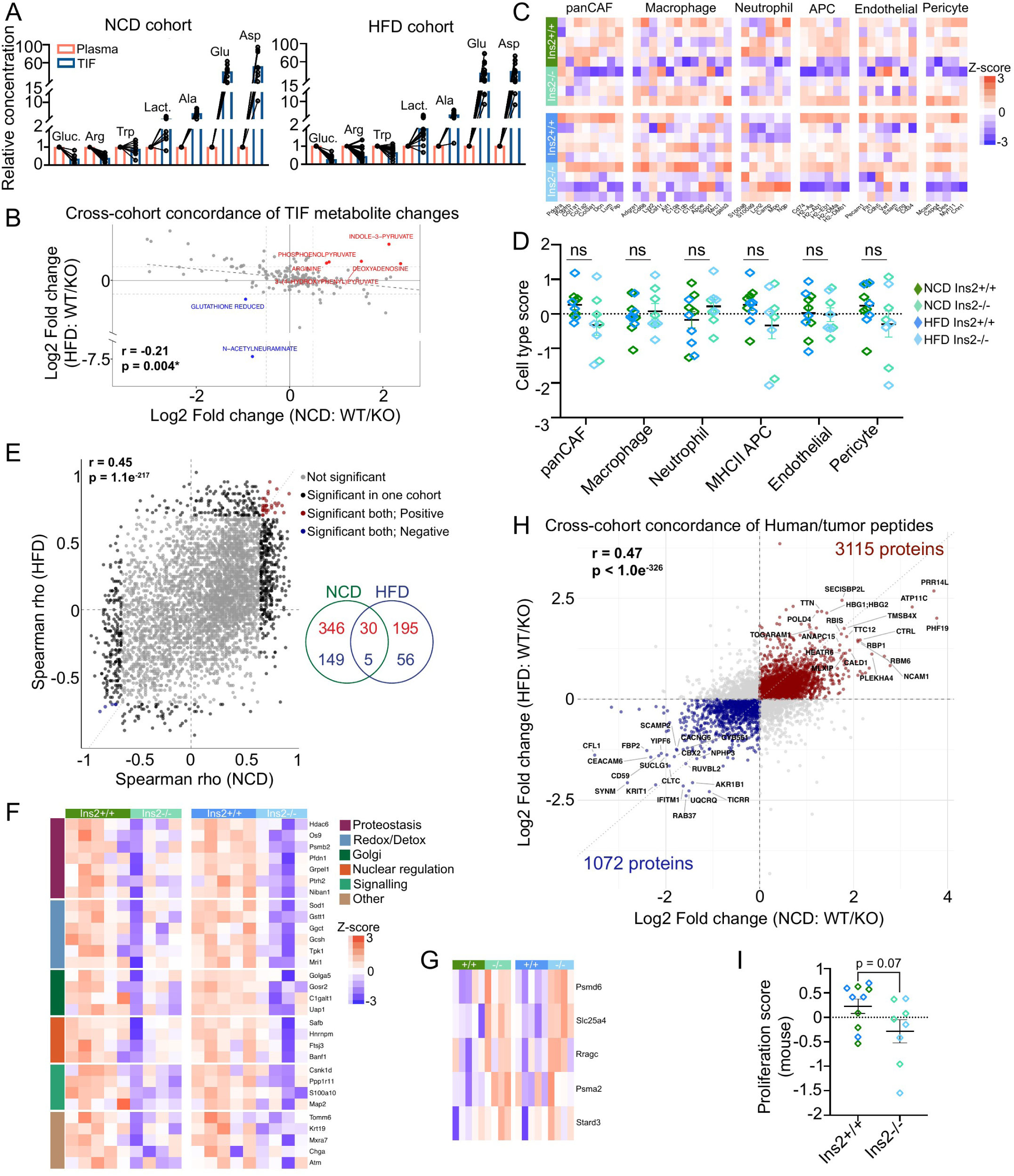
Metabolomic and proteomic characterization of PDAC from male mice, related to Figures 6 and 7. (A) Concentration of metabolites in tumor interstitial fluid (TIF) relative to the matched plasma values of samples from chow-fed (left) and HFD-fed (right) male cohorts. (B) Cross-cohort concordance analysis of relative changes in metabolite levels between genotypes in TIF samples from chow-fed and HFD-fed male cohorts. r and p values were determined by Pearson correlation analysis. (C-D) Heatmap showing the relative abundance of cell-type markers (mouse-specific proteins) across common stromal cell types in PDAC (C), and the quantification of the mean abundance of cell-type markers between genotypes (D). ns p > 0.05 by one-way ANOVA. (E) Concordance analysis of the Spearman correlation between each mouse/stromal protein and the corresponding insulin score in both diet conditions. (F-G) Heatmaps that summarize the proteins showing significant positive (F) and negative (G) correlation with the host’s insulin score from the concordance correlation analysis (E). (H) Scatter plot comparing the effect of *Ins2* deletion on human/tumor-specific proteome in tumors from the chow-fed and HFD-fed cohorts. Proteins showing concordant increases in tumors from *Ins2^+/+^* hosts in both cohorts are highlighted in red. Proteins showing concordant decreases are highlighted in blue. All other proteins are shown in grey. r and p values were determined by Pearson correlation analysis.

**Figure S7.**
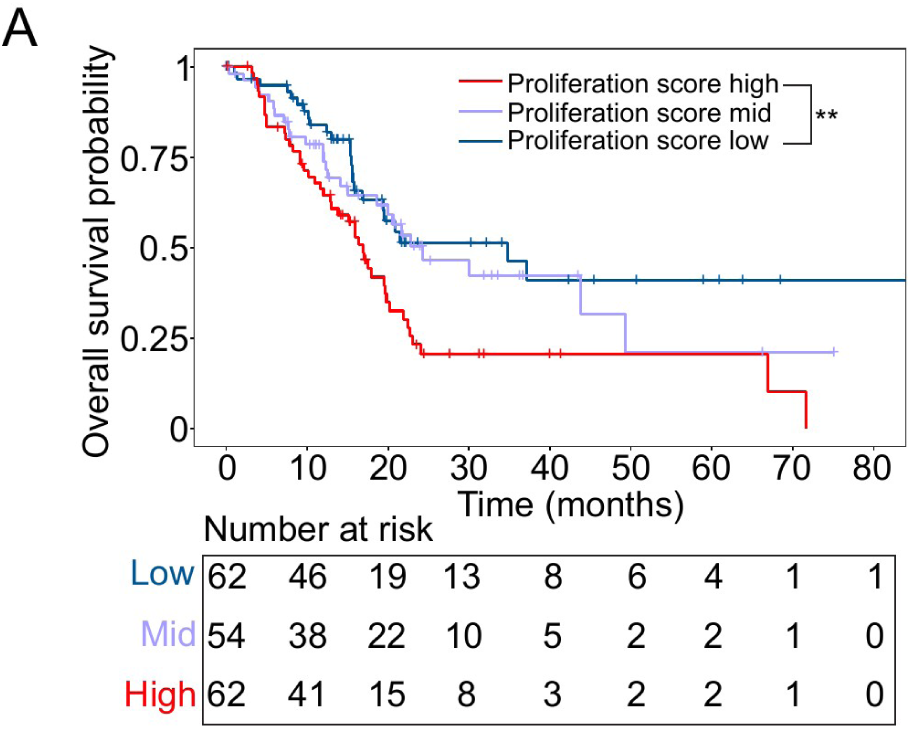
Higher expression of proliferation marker genes is associated with worse patient survival, related to Figures 7. (A) Kaplan-Meier survival analysis of patient (TCGA PAAD) tumors with high and low mean expression of the proliferation markers in Fig. 6G. **p < 0.01 by log-rank test.

